# Viral proteogenomic and expression profiling during fully productive replication of a skin-tropic herpesvirus in the natural host

**DOI:** 10.1101/2023.02.13.528429

**Authors:** Jeremy Volkening, Stephen J. Spatz, Nagendraprabhu Ponnuraj, Haji Akbar, Justine V. Arrington, Keith W. Jarosinski

## Abstract

Efficient transmission of herpesviruses is essential for dissemination in host populations; however, little is known about the viral genes that mediate transmission, mostly due to their close relationship to their natural host. Marek’s disease is a devastating herpesviral disease of chickens caused by Marek’s disease virus (MDV) and an excellent natural model to study skin-tropic herpesviruses and transmission. Similar to varicella zoster virus that causes chicken pox in humans, the only site where fully productive replication occurs is in epithelial skin cells and this is required for host to host transmission. Here, we enriched for actively replicating virus in feather follicle epithelial skin cells of live chickens to measure both viral transcription and protein expression using combined RNA sequencing and LC/MS-MS bottom-up proteomics. Enrichment produced a previously unseen breadth and depth of viral peptide sequencing. We confirmed protein translation for 84 viral genes at high confidence (1% FDR) and correlated relative protein abundance with RNA expression levels. Using a proteogenomic approach, we confirmed translation of most well-characterized spliced viral transcripts and identified a novel, abundant isoform of the 14 kDa transcript family via both intron-spanning sequencing reads as well as a high-quality junction-spanning peptide identification. We identified peptides representing alternative start codon usage in several genes and putative novel microORFs at the 5’ ends of two core herpesviral genes, pUL47 and ICP4, along with evidence of transcription and translation of the capsid scaffold protein pUL26.5. Using a natural animal host model system to examine viral gene expression provides a robust, efficient, and meaningful way of validating results gathered from cell culture systems.

**Author Summary:** In the natural host, the transcriptome and proteome of many herpesviruses are poorly defined. Here, we evaluated the viral transcriptome and proteome in feather follicle epithelial skin cells of chickens infected with Marek’s disease virus (MDV), an important poultry pathogen as well as an excellent model for skin-tropic human alphaherpesvirus replication in skin cells. Using fluorescently tagged virus, we significantly enriched the number of infected cells sampled from live chickens, greatly enhancing the detection of viral transcripts and proteins within a host background. Based on this, we could confirm the translation of most transcripts using deep MS/MS-based proteomics and identify novel expressed peptides supportive of an increasingly complex translational and regulatory viral landscape. The demonstrated deep peptide sequencing capability can serve as a template for future work in herpesviral proteomics.

## Introduction

Herpesviruses have two modes of spread, cell-to-cell and cell-free virion release, where the significant advantage of cell-to-cell spread is the evasion of the immune system. However, infectious cell-free virus must be released into the environment to disseminate amongst a population [1]. Herpesviruses are highly adapted to their host species, having co-evolved for millions of years, which makes studying natural virus transmission in the host population difficult, and for humans nearly impossible. In addition, most herpesviruses are primarily cell-associated in cell culture and within the host, where virions are delivered through cell-cell junctions and tunneling nanotubes [2]. Depending on the herpesvirus, infectious cell-free virus released from cells in cell culture is highly variable. Most studies have focused on in vitro cell culture models to study herpesvirus replication that is primarily cell-to-cell spread. As the transcriptional and translational machinery active during the cell-associated and cell-free (fully productive) stages of the viral life cycle is likely to vary significantly, we sought to address fully productive virus replication using a natural herpesvirus animal model system.

Marek’s disease virus (MDV; *Gallid alphaherpesvirus 2*; GaHV3) is a significant pathogen affecting the poultry industry due to its global distribution and transmissibility. MDV is a member of the *Herpesviridae*, subfamily *Alphaherpesvirinae*, and is related to the human herpes simplex virus 1 (HSV-1), HSV-2, and varicella-zoster virus (VZV), both genetically [3, 4] and, in particular, in their shared tropism to epithelial skin cells required for replication, egress, and dissemination into the environment. In contrast to most other alphaherpesviruses, MDV is strictly cell-associated when grown in cell culture through semi-productive replication, relying on spread through cell-to-cell contact. To date, no cell-free virions have been produced using primary cell culture or engineered immortalized continuous cell lines [5–10]. The only cells known to facilitate cell-free virus release, or fully productive virus replication, are differentiated chicken epithelial skin cells called the feather follicle epithelium [11]. The production of cell-free virus is required for host-to-host transmission, and specific viral genes required for this process have been identified [12]. The related human VZV is also primarily cell-associated in cell culture, with only small amounts of infectious cell-free virus produced, while the prototype alphaherpesvirus HSV-1 generates cell-free virus that is partially dependent on the cell type used for infection. However, for these skin-tropic viruses, human-to-human transmission cannot be studied, and mouse models do not facilitate transmission. Thus, the MDV-chicken model system is well suited to address transmission and fully productive replication in the host.

The ∼180 kb double-stranded DNA genome of MDV was first sequenced in 2000 for the very virulent Md5 [13] and attenuated GA [14] strains, annotated to have 338 open reading frames (ORFs) of >60 aa in length, of which 103 ORFs were predicted to be functional. The current annotation of the MDV genome largely relies on both *in silico* ORF predictions and homologous ORFs in related alphaherpesviruses [15]. Recently, studies on the MDV transcriptome have been reported in cell culture [16] and *in vitro* infected B cells [17], expanding our knowledge of MDV’s complex gene expression patterns in different types of cells. In addition, the transcriptome of MDV-infected feathers in chickens has been recently reported, with limited success in identifying viral transcripts deemed necessary for productive infection [18]. Mass spectrometry (MS)-based proteomics studies have also provided useful information on viral proteins produced in MDV-transformed chicken cells [19] and during *in vitro* replication in cell culture [20] and B cells [17]. Together, these studies have provided a foundational understanding of viral transcription and translation, but they are limited either by an *in vitro* context or a shallow breadth of coverage depth.

Over the past decade, we have established a robust natural infection system by which we can identify and enrich MDV-infected epithelial skin cells from live chickens using fluorescence microscopy without complex manipulation of the samples [21–23]. To further our knowledge of the viral machinery active during the critical stages of virus assembly, egress, and shedding, we herein combined this system with RNA sequencing and bottom-up proteomics to define the combined viral transcriptome and translated proteome during fully productive replication within the natural host.

## Results & Discussion

### Visualization of the transcriptional and translational landscape in epithelial skin cells

The overall experimental design is shown in Figure 1 and described in the Materials and Methods. Briefly, uninfected and MDV-infected samples were processed for RNA sequencing and LC-MS/MS analyses. Overviews of strand-specific read coverage detected for RNA-Seq-based splicing events and MS/MS peptides are shown for the unique long (S1 Fig), repeat long (S2 Fig), repeat short (S3 Fig), and unique short (S4 Fig) regions. These genome tracks and additional data tracks generated from this study can also be viewed interactively at https://igv.base2.bio/AAG3-9Fja-99a2-2asZ/.

**Fig 1.**
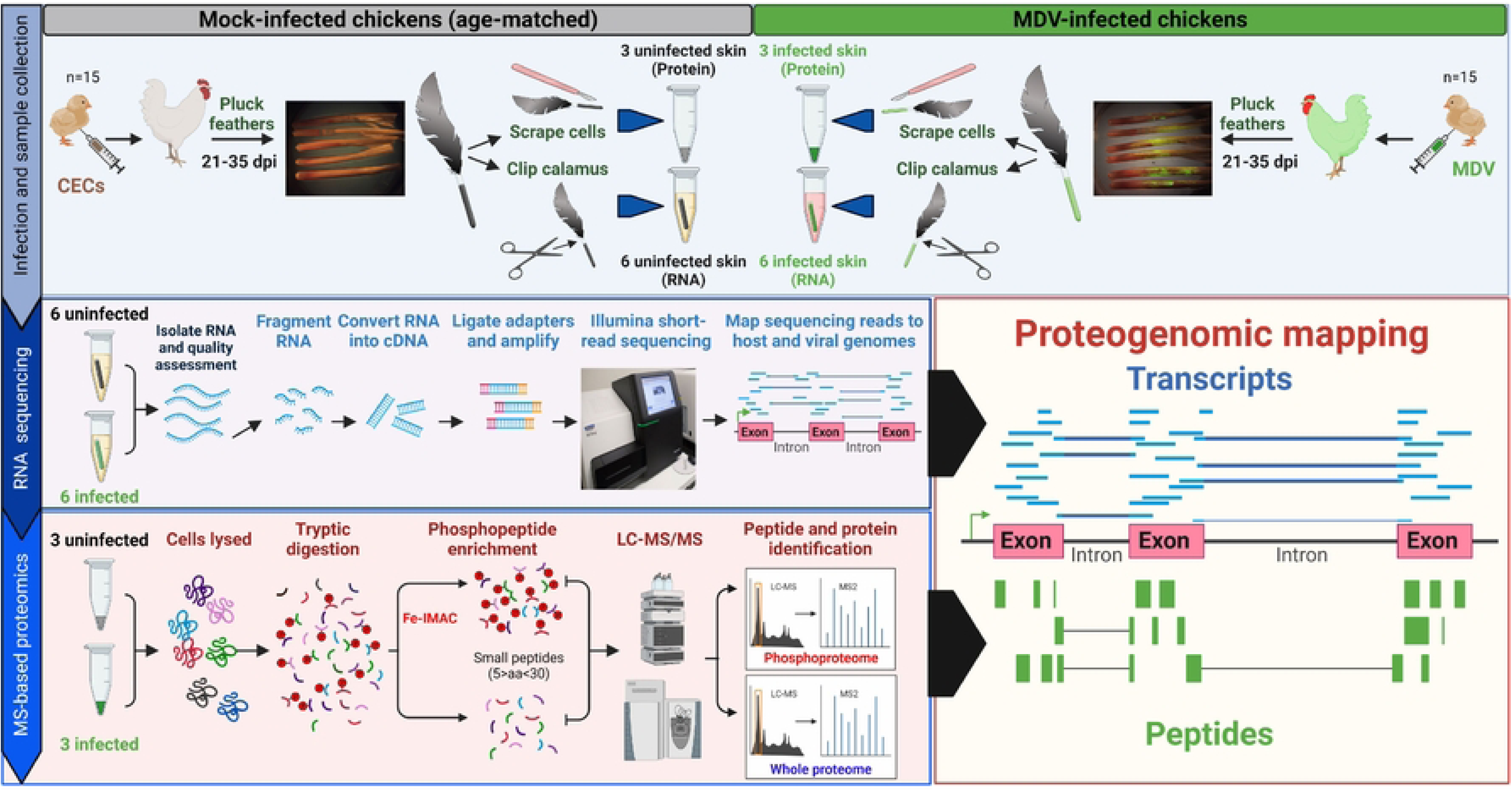
Schematic illustration of the experimental approach. Day old chicks were experimentally infected with MDV or left uninfected. Feathers were plucked weekly and birds heavily infected in feathers were sacrificed for sample collection, along with an equal number of age-matched controls. Samples were collected between 21 and 35 days based on the level of pUL47eGFP. Approximately 6-10 feathers per bird (n=6/group) were directly clipped at the calamus and dropped in ice-cold RNA STAT60 and snap frozen on dry ice then stored at -80°C until processed for RNA extraction and subsequently RNA sequencing (see Materials and Methods). Approximately 3-4 feathers per bird (n=3/group) were used to collect protein by scraping fluorescent cells into an Eppendorf tube and all samples stored at -80°C until processed for MS-based proteomics (see Materials and Methods).

### The viral proteome during fully productive replication

LC-MS/MS-based bottom-up proteomics was used to examine the expressed proteome of MDV in epithelial skin cells. Both total protein and phospho-enriched samples were used to increase the coverage of total viral proteins (Fig 1C). Each of the six replicates (3 infected, 3 uninfected) produced 105k-122k total MS2 spectra from the unenriched samples and 68k-88k MS2 spectra from the phospho-enriched fractions (S1 Table). Spectra matched to peptides (peptide-spectrum matches; PSMs) ranged from 13k-23k for unenriched samples and 4k-11k for phospho-enriched samples [1% PSM false discovery rate (FDR)]. Of these PSMs, the fraction matching to MDV proteins ranged from 5.6-6.1% for infected unenriched samples and 8.7-9.4% for infected phospho-enriched samples. Total MDV-matched PSM counts in uninfected replicates were 0 or 1 for both unenriched and phospho-enriched fractions, suggestive of a very high specificity in peptide assignment. The MDV PSMs represented 1,484 distinct annotated viral peptides identified at 1% peptide FDR, not including different isoforms of each peptide such as post-translational modifications (PTMs) and missed cleavages, which are listed in S2 Table.

A total of 84 non-redundant MDV proteins (excluding terminal repeat copies but including different splice forms) were identified by at least one peptide at a maximum protein q-value of 0.01 (S3 Table). Of these, 79 proteins were identified by at least two distinct peptides at 1% FDR (commonly used criteria for protein presence), and 80 of the 84 proteins were detected in all three biological replicates. When considering expected peptide coverage (defined here as the protein length covered by detected peptides as a fraction of the total residues found in theoretical tryptic peptides ≥ 6 aa), 47 proteins had a breadth of coverage > 50%, 18 proteins had coverage > 80%, and five proteins were over 90% covered. Histograms of unique peptide counts and breadth of coverage for detected viral proteins are shown in S5 Fig. Based on relative iBAQ (sum of peptide precursor intensities divided by theoretically observable peptides and divided again by the sum of all iBAQs), the most abundant protein present in epithelia skin cells was glycoprotein C (gC), followed by UL45 (envelope protein), UL42 (DNA Pol accessory), UL39 and UL40 (ribonucleotide reductase subunits), UL50 (Deoxyuridine 5’-triphosphate nucleotidohydrolase; DUT), UL49 (tegument protein VP22), and UL18 and UL19 (capsid subunits) (S3 Table). These nine proteins comprised an estimated 76% of the quantified viral protein load by molarity.

#### Single-peptide proteins

Using a single peptide as evidence of protein expression is generally unreliable, as even high-scoring PSMs can be incorrect due to factors such as an incomplete search database or unsearched PTMs. However, manual analysis can help to provide additional support either for or against the peptide match and confidence in the presence of the matched protein. Here, we used visual inspection of the matching MS2 b/y ion series and the extracted ion chromatogram (XIC) for the peptide mass and retention time to further evaluate peptides from single-peptide proteins. The use of the XIC was informative here because no elution peak would be expected in uninfected replicates.

Protein UL11 (pUL11-CEP3) was identified from a single peptide in both phosphorylated and unphosphorylated forms in all three infected replicates. pUL11 is a short protein with only five predicted tryptic peptides ≥ 6 aa. Inspection of the MS2 spectra from three selected top PSMs of the unphosphorylated peptide shows a nearly complete y-ion series within the detectable m/z range and with no precursor co-isolation (S6 Fig). Additionally, elution peaks in the XIC plot are found only for the three infected replicates. This supplementary evidence provides strong support for identifying this peptide and thus, the expression of pUL11 in the skin cells during productive replication. A single (different) peptide from pUL11 was formerly identified in MDV-infected CECs [20], and it should be noted that most studies using MS-based proteomics do not typically detect this protein with efficiency [17, 24, 25]. For pUL49.5 or glycoprotein N (MDV064), inspecting the MS2 spectra and XIC for the single identified peptide provides similarly strong support for its identification and presence (S7 Fig). However, an inspection of the MS2 spectra and XICs for the remaining three single-peptide proteins (MDV076/MEQ: SHDIPNSPS[+80]K; MDV093/SORF4: SRDFS[+80]WQNLNSHGNSGLR; MDV091.5: TINESLVPANPVPRT[+80]PVPSGGFVLTIGR), are less convincing with less-complete b/y ion series and noisier XIC peaks (S8 Fig) that neither strongly support nor dispute the identifications.

### The MDV transcriptome of epithelial skin cells

A summary of RNA-Seq data for this experiment is shown in S4 Table. Library sizes for the twelve replicates ranged from 9.4×10^6^ to 1.3×10^7^ read pairs after trimming/filtering. The fraction of reads mapping to the MDV (GaHV2) genome for infected samples ranged from 9.3% to 24.6%. For uninfected replicates, the fraction of MDV-mapped reads was negligible (from 0 to 26 *total* mapped read pairs) – as with MS/MS, identification of MDV reads was highly specific to infected samples.

The library’s overall strand specificity (number of read pairs mapping to the expected strand as a fraction of all classified read pairs) was high (range 95-98%). However, it was observed that the strand specificity of viral reads was significantly lower (range 66-75%). For reference, the expected specificity for randomly distributed (i.e., not strand-specific) libraries would be 50%. From these values and visual inspection of the read alignments in IGV, there appeared to be a significant fraction of viral reads mapping to both presumed intergenic and presumed anti-sense regions of the viral genome. Visualization of host read alignments, on the other hand, appeared highly strand-specific and highly intragenic (Volkening, unpublished observation). Because of this apparent background noise, possibly due to viral gDNA contamination, it was decided that traditional count-based or k-mer based methods of gene expression analysis would be unsuitable. Instead, the median read depth was calculated for each viral gene interval and replicate, as described in the Materials and Methods, and summarized to an average median read depth and standard deviation across all six replicates. Similarly, a baseline read depth distribution was calculated for intergenic/antisense regions (S9 Fig). A median depth threshold of two standard deviations above the mean background coverage (50-fold + 2 × 33-fold = 116-fold) was then used as the threshold for the “expression” of a gene with ∼98% one-tailed confidence.

Approximately 75% (114/152) of non-redundant viral genes found in the RB-1B annotations used here were detected as expressed at the mRNA level above this threshold. Of the 152 gene models in the annotations used herein, 55 are annotated as “hypothetical proteins,” likely based on *in silico* ORF prediction alone, and many overlap core or well-characterized genes on the strand. When these are ignored, we find evidence for the expression of 89/97 (92%) of the remaining annotated non-redundant genes in epithelial skin cells.

The most highly expressed transcript was MDV075.1/B68 (S3 Table). However, there was no evidence of the translation of this coding sequence in the MS/MS dataset. The expression values are likely a result of non-spliced mRNA from the overlapping, and highly expressed, 14 kDa protein gene family. B68 lies within the intron of MDV075 (14 kDa A), downstream of the first intron donor site (Fig 2). Of note, the next most highly expressed gene was MDV082 (S3 Table), located in the short-inverted repeat (IRS) downstream of ICP4. Little was known about this gene until recently where it was found to be expressed late in the viral life cycle and enhances the rate of disease progression, but it was not essential for replication, spread, or tumor formation [26]. We also found evidence of abundant expression of MDV082 at the protein level (∼ 1% relative molar abundance, 92% protein coverage). MDV057 (UL44-gC) transcripts were similarly highly abundant, mirroring their high abundance at the protein level.

**Fig 2.**
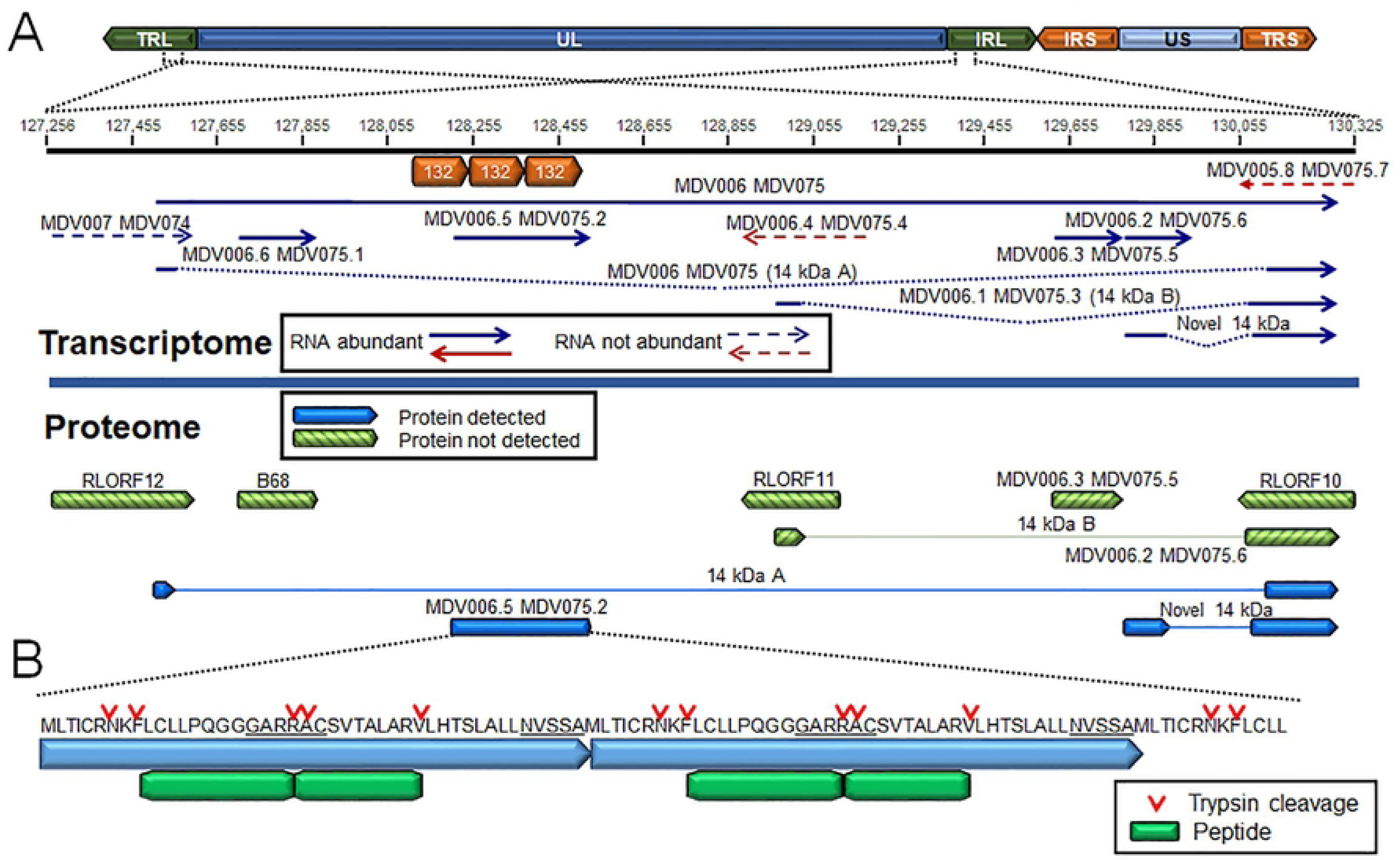
Expression of the 1.8 kb family transcripts, novel mRNA splicing, and validation of protein expression in epithelial skin cells. (A) Schematic representation of the MDV genome and location of the terminal (TRL) and internal (IRL) repeat long, unique long (UL), terminal (TRS) and internal (IRS) repeat short, and unique short (US) regions. The region encoding the 1.8 kb family transcripts is expanded from the TRL and IRL. A summary of transcripts detected or not detected in RNA sequencing and proteomics are shown along with the region of the 132 bp direct repeats. (B) Validation of the 132 bp direct repeat ORF encoding the MDV006.5 MDV075.2 protein. Peptides detected are noted in the figure legend along with tryptic cleavage sites.

Of the annotated coding sequences with read depths below the threshold level, most (30/38) are ORFs annotated as “hypothetical protein” and are unlikely to be functional in epithelial skin cells, including MDV013.5 (LORF4), MDV057.8 (LORF8; 23 kDa protein), MDV074 (RLORF12), MDV075.7 (RLORF10), and MDV077 (23 kDa nuclear protein). Of these genes, only MDV072 (LORF5) has evidence of expression at the protein level (four distinct peptides observed, q = 0.0004). Core herpesviral genes in this category included MDV017 (UL5) and MDV066 (UL52) which are both helicase-primase subunits. Both were detected with high confidence at the protein level, but, along with LORF5, they are the least abundant proteins present by estimated molarity (riBAQ) (S3 Table).

Overall, the correlation between RNA and protein levels was moderate (Pearson R for log2 median read coverage vs. log2 riBAQ protein abundance = 0.62) (S10 Fig). However, similarly low levels of RNA/protein correlation are frequently observed due to differences in transcript vs. protein stability as well as measurement error.

### mRNA splicing in MDV

Novel mRNA splicing has been identified during *in vitro* infection of CECs [16, 27] and, more recently, B cells [17], in particular MDV073 (pp38), MDV078 (vCXCL13-vIL8), and MDV027 (UL15-TRM3). As in other transcriptomic studies with MDV-infected cells [16, 17], we also detected mRNA splicing events occurring in known coding regions and seemingly intergenic and anti-sense regions (S1-S4 Tables). Reads spanning some of these introns were detected in our RNA-Seq analysis but at very low abundance, suggesting their expression is limited in epithelial skin cells. Similarly, formerly identified mRNA splicing of MDV076 (Meq) and MDV078.3 (RLORF4) to MDV078 (vCXCL13-vIL8) were detected [28–30], but also at low levels in epithelial skin cells. In contrast to the above splice variants detected at low abundance, pp38A and pp38B transcripts were not detected (S1 Fig), suggesting they are not expressed in epithelial skin cells. In immortalized chicken cells (DF-1), primary chicken cells (CEC and CKC), and splenocytes infected with MDV, the expression of both pp38 and pp38B at the RNA level has been demonstrated, although no proteomic evidence (MS or western blotting) was reported [27]. We identified a novel pp38 splice variant (Novel pp38C) at the RNA level, albeit without evidence from MS/MS discussed below. Only minor splicing events of pp24 were detected in our RNA-Seq data, contrary to what was previously reported by Bertzbach et al. [17]. Overall, it appears increasingly likely that the extent of viral mRNA splicing identified within the IRL/IRS regions may depend on the infected cell type. Viral gene expression is likely more tightly regulated in cells naturally infected by MDV compared to artificial *in vitro* cell culture systems that do not facilitate fully productive replication.

### Confirmation of productive transcript splicing by MS/MS

Decades of gene expression and mRNA splicing studies have identified numerous viral mRNA splicing events during cell culture replication and in MDV-transformed chicken cells, including vIL-8, Meq/vIL-8, pp38, RLORF4, and gC splicing products [22, 27, 28, 31–33]. Although extensive analysis has been performed at the transcriptional level using RT-PCR and sequencing, in addition to traditional protein expression studies using western blotting and immunofluorescence assays, few of these spliced products have been directly validated at the level of peptide identification. Due to the depth of peptide sequencing achieved herein, we could detect high-confidence peptides spanning the previously described intron boundaries of four proteins. These include the second intron of vCXCL13 (RTEIIFALK), UL15 (STVTFASSHNTNSIR), gC104 (DGSLPDHRS[+80]P), and the 14 kDa A nuclear protein (YISYPGCIDCGPTFHLETDTATTR) (Fig 3). Three of these peptide identifications, in addition to q-values < 0.01, are supported by rich y/b ion series and infection-specific elution profiles (S11 Fig). The pUL15 peptide was found only by the Byonic search engine, and the supporting MS2 and XIC plots are poor (S12A Fig). Additional novel splice junctions are described as part of the proteogenomic search results below.

**Fig 3.**
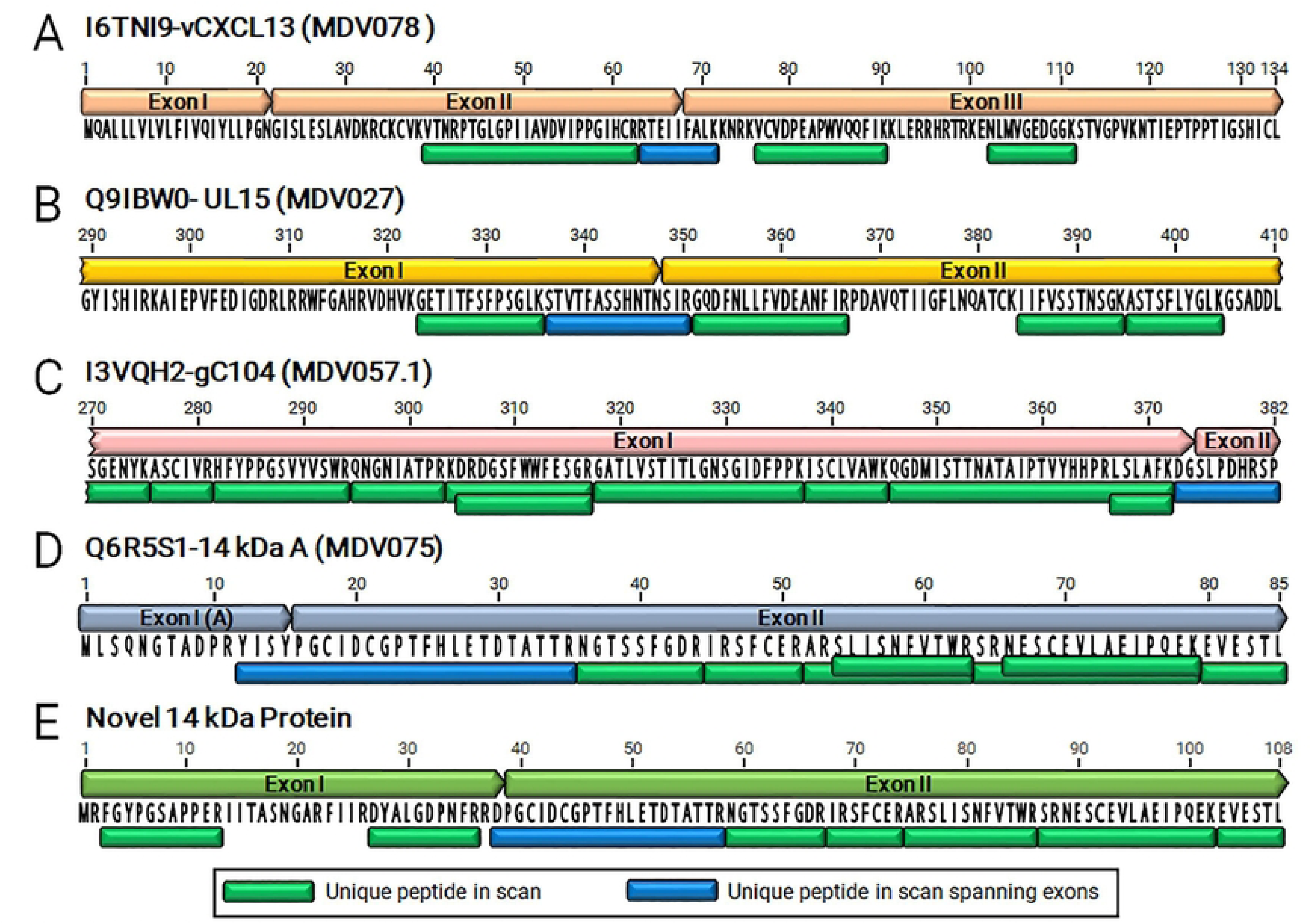
Evidence of mRNA splice products by unique peptides. Following MS/MS-based proteomics and analyses, unique peptides spanning exon junctions were identified for vCXCL13- vIL8 (A), pUL15 (B), gC104 (C), 14 kDa A (D), and the newly identified Novel 14 kDa protein (E). Peptides detected using tryptic digestion are shown with peptides spanning exon junctions in dark blue.

### Validation of translation start sites and assessment of protein N-terminal PTMs

MS/MS database searching of enzymatic digest spectra allows for the identification of protein N-terminal peptides, which lack a cleavage site (in this case, by trypsin/LysC) at the N-terminal end of the peptide. By including the common protein N-terminal PTMs of N-terminal methionine excision (NME) and N-terminal acetylation (NTA) in database searches, we confirmed the translational initiation sites (TIS), as well as PTM state, for 23 viral proteins (S5 Table). Nearly all identified TISs agreed with the existing gene annotations. By including the sequences for potential alternative TISs (based on the start codon and Kozak context) in the search database [34], we could also search for alternative or incorrectly annotated sites. For MDV015 (UL3), we confirmed the use of a downstream start codon as the TIS, in agreement with the RefSeq annotation of Md5 (NC_002229). Although there is an in-frame start codon upstream to this position, it has a weak Kozak context (CAC**ATG**C) compared to the strong Kozak consensus of the true TIS (ATA**ATG**G). For this analysis, a strong Kozak motif was considered to be one matching the consensus sequence (-3)RNN**ATG**G(+4). In addition, we identified a peptide indicating an alternative TIS for MDV055 (UL42), together with a peptide representing the annotated TIS (S13 Fig). The annotated TIS peptide (AGITMGSEHMYDDTTFPTNDPESSWK) was identified by a total of 23 PSMs in all three replicates and in two different charge states (S13B Fig). The alternative TIS peptide (GSEHMYDDTTFPTNDPESSWK) was identified by a total of 12 PSMs in all three replicates and in two different charge states (S13C Fig). Similarly, normalized LFQ intensities for the two peptides were 1.1×10^7^ ± 4.2×10^6^ and 2.4×10^6^ ± 2.5×10^5^ (S6 Table), suggesting the alternative downstream TIS is utilized at roughly one-fourth the rate of the annotated TIS. Both N-terminal peptides underwent NME and NTA. The alternative TIS (S13A Fig) has a slightly stronger Kozak context, with conserved bases at both -3 and +4 positions (ACT**ATG**G), while the annotated TIS has a non-conserved -3 base (TCA**ATG**G). Both peptides have strong MS2 ion series and infection-specific extracted ion chromatograms. The biological significance of the alternative TIS of MDV055 (UL42) in epithelial skin cells remains unknown.

Several additional novel N-terminal peptides were detected in the expanded search. A putative alternative TIS peptide ([+42]S[+80]SSTLAQIPNVYQVIDPLAIDTSSTSTK) was found for MDV070 (UL55) in addition to detecting the annotated TIS peptide ([+42]AAGAMS[+80]SSTLAQIPNVYQVIDPLAIDTSSTSTK). In this case, only the alternative TIS (S14A Fig) has a strong Kozak context (GCG**ATG**T). Both peptides have sparse but specific MS2 ion series, and both have extracted ion chromatograms specific to infected replicates (S14BC Fig). The annotated TIS peptide was identified in four PSMs from all three phospho-enriched replicates, while the alternative TIS peptide was identified in two PSMs from two phospho-enriched replicates. Both N-terminal peptides underwent NME and NTA (S5 Table).

Detection of a novel N-terminal peptide (ANINHIDVPAGHSATTTIPR) in MDV096 (US7-gE) would represent translation initiation at a non-canonical start codon in an otherwise strong Kozak context (GGA**ACG**G). The peptide was identified in all three replicates and with a reasonably complete ion series; however, the extracted ion chromatogram shows elution peaks in both infected and uninfected samples, suggesting that this is likely a misidentified peptide (S12D Fig).

Overall, patterns of NME and NTA followed the expected rules based on the local amino acid context (S5 Table). Of the 23 viral protein N-terminal peptides detected in MS/MS (including alternate starts), 15 were always detected with the N-terminal Met removed. All of these had small +2 amino acids (A, G, S, or T) in accordance with known rules [35]. Six TIS-indicating peptides always retained their N-terminal Met. All of these had larger penultimate residues (D, E, or M). Two termini were identified with peptides both N-terminally cleaved and uncleaved, somewhat surprisingly (M and N +2 residues). Nearly all identified N-terminal peptides were acetylated, either in every PSM (19) or part of the time (3). Only one peptide was detected in an unacetylated state - the N-terminus of MDV044 (UL31), TGHTLVR.

### Quantitative analysis of mRNA splicing and validation at the protein level in epithelial skin cells

The alphaherpesvirus conserved gC, encoded by MDV057 or UL44, has been previously shown to be alternatively spliced to produce two secreted forms called MDV057.1 (gC104) and MDV057.2 (gC145) [22]. The mRNA splicing of UL44 is believed to be conserved, as it has been observed during HSV infection [36] as well as in turkey herpesvirus (HVT) and *Gallid alphaherpesvirus* 3 (GaHV3) replication (Jarosinski, unpublished data). Importantly, all three MDV gC proteins (gC, gC104, and gC145) are required for efficient horizontal transmission [22], but their expression at the mRNA or protein level had never been examined in epithelia skin cells. All three forms have been detected in RNA-Seq studies in infected cell cultures [16, 22] and in B cells infected *in vitro* [17]. Here, we confirmed that both splice variants are expressed in epithelial skin cells with gC104 (MDV057.1) ∼4-fold more abundant than gC145 (MDV057.2) and ∼10-fold more abundant than the non-spliced transcript based on intron read depth (Fig 4A). Five peptides that are unique to the non-spliced product (MDV057) were detected confirming full-length gC expression (Fig 4B). As discussed above, the unique intron-spanning peptide of gC104 was detected, while no unique peptides were identified for gC145; however, the transcript abundance data suggests it is present, and most peptides would be shared with the other isoforms. Tryptic mapping of gC145 showed predicted gC145 unique fragments of 3, 4, 1, and 38 aa (Fig 4C). Only the 38 aa peptide would be detectable by MS/MS – it is therefore entirely possible that this proteoform is present but that the single unique peptide was missed. Further studies are needed to determine where UL44-gC145 is expressed at the protein level (gC145). However, we can confirm gC104 is expressed at both the RNA and protein levels.

**Fig 4.**
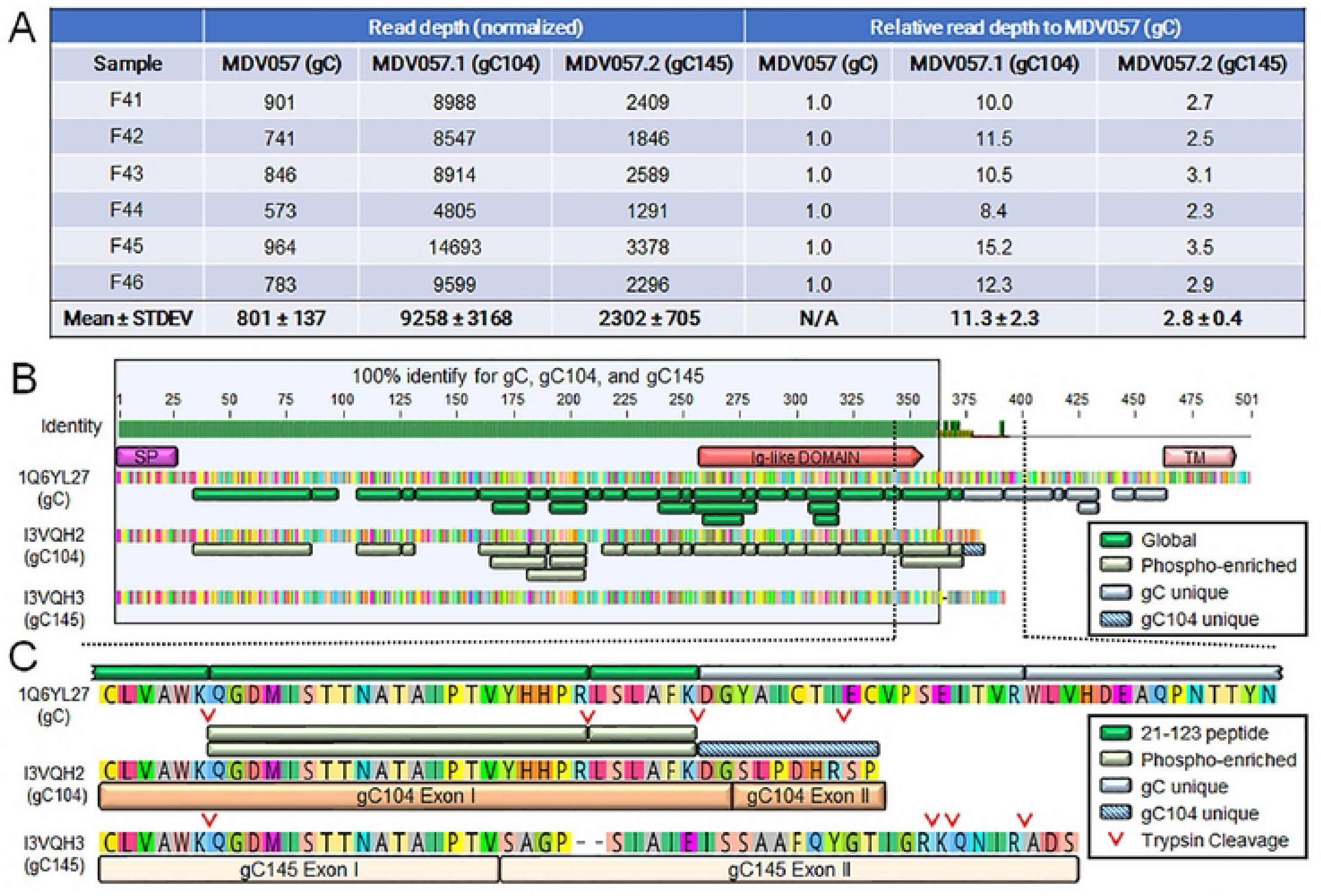
Quantitative analysis MDV057, MDV057.1, and MDV057.2 mRNA expression and peptide validation. (A) Total mRNA reads for MDV057 (gC), MDV057.1 (gC104), and MDV057.2 (gC145) in the six infected replicates and the average reads ± standard deviations are shown in table form. The ratio of MDV0057.1 (gC145) and MDV057.2 (gC145) transcripts compared to MDV057 (gC) ± standard deviations is also shown. (B) Protein alignment of gC, gC104, and gC145 using MUSCLE alignment. Also shown are the predicted signal peptide (SP), Ig-like, and transmembrane (TM) domains predicted using SignalP-6.0 [80], DeepTMHMM [81] and MyHits [82]. Unique peptides were assigned to IQ6YL27 (gC) in the global protein analysis (21-123) and to I3VQH2 (gC104) in phospho-enriched samples with one peptide specific for gC104. (C) The region highlighted in (B) was expanded to show the protein sequences of gC, gC104, and gC145, exon junctions for gC104 and gC145, and unique peptides detected for gC and gC104. Predicted tryptic cleavage sites are also shown for gC104 and gC145.

The MDV-specific pp24 and pp38 phosphoproteins are encoded by MDV008 and MDV073, respectively, and share identical N-terminal 65 aa sequences (S15 Fig). Additionally, the MDV073 gene has been shown to produce two alternatively spliced mRNA and proteins called spl A and spl B [27], or pp38A and pp38B, respectively, during *in vitro* replication, and they are reported to be important for MD pathogenesis and tumor development [37]. Analysis of intron-spanning reads for this region showed no evidence that pp38A or pp38B were produced in epithelial skin cells; however, a novel splicing product was identified utilizing the pp38A exon I donor splicing site (D1) to a novel acceptor site (A) and spanned by an average of 114±56 reads in the six replicates. This acceptor site was located downstream of the pp38 ORF stop codon and would code for a protein of 137 aa that we termed Novel pp38C, which only differs from pp38A by the C-terminal 8 aa (S15A Fig).

### RNA expression and protein validation of pUL26.5 in epithelial skin cells

The overlapping ORFs encoding MDV038 (UL26) and MDV039 (UL26.5) pose a differentiation dilemma. The UL26 and UL26.5 proteins are encoded by overlapping transcripts with alternative TIS, with the encoded proteins sharing identical C-termini. Therefore, nearly all the potential peptides of the latter are shared with the former (Fig 5A). UL26 encodes a serine protease (pUL26-SCAF) cleaved during procapsid maturation to yield two proteins, VP24 and VP21, the latter of which is almost identical to pUL26.5-ICP35 [38–43]. Fortuitously, a tryptic peptide of the N-terminus of UL26.5 was detected in our data with strong elution peaks in infected replicates (Fig 5B). This distinguishable finding, along with the increased RNA-Seq depth (Fig 5C) and more intense LFQ peptide intensities for UL26.5 vs. the full UL26 (S6 Table), strongly suggests pUL26.5-ICP35 is produced from its own TIS in epithelial skin cells. The Neural Network Promoter Prediction program [44] suggests two potential promoters at ∼100 and ∼250 bp upstream of the pUL26.5 TIS (Fig. 5D). A similar genetic arrangement with an internal promoter within the body of UL26 has been reported for HSV-1, suggestive of a conserved transcriptional regulatory function [45].

**Fig 5.**
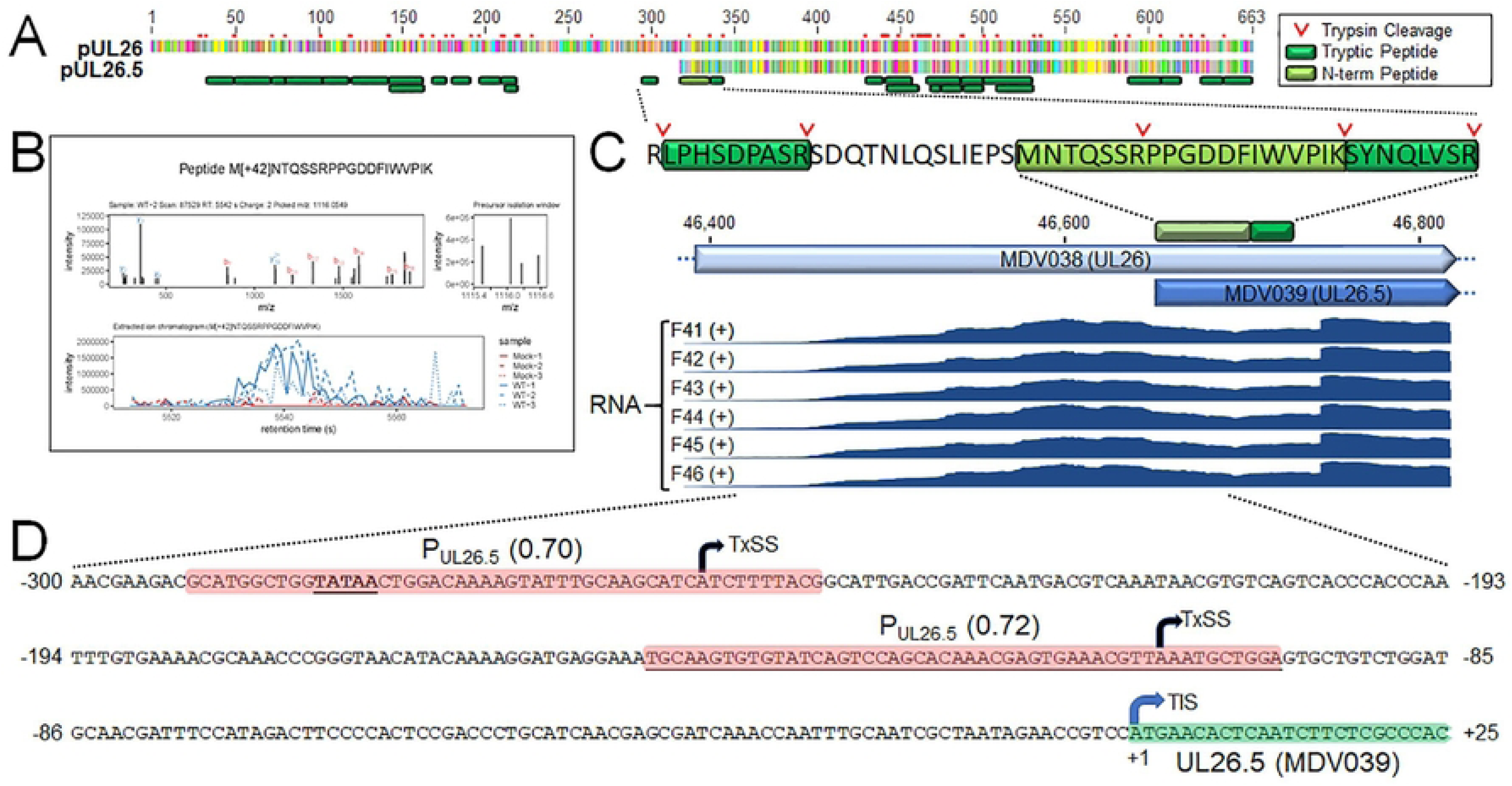
Expression of pUL26.5 in epithelial skin cells. (A) MUSLE alignment of pUL26 (Q19BC6) and pUL26.5 (A0A2H4V874) and peptides detected in MS/MS. Tryptic cleavage sites are shown. (B) The ion series and extracted ion chromatogram of the N-terminus of UL26.5. (C) The N-terminal peptide of pUL26.5 detected in MS/MS with tryptic cleavage sites are shown. RNA sequencing reads for the six replicates are shown below showing increased transcription upstream of MDV039 (UL26.5). (D). The Neural Network Promoter Prediction program predicted two putative promoters at ∼250 and ∼100 bp from the TIS with potential transcription start sites (TxSS) with 0.70 and 0.72 scores of predictability.

### Evidence for translation of the 132 bp direct repeat reading frame

The hypothetical MDV075.2 ORF is highly variable in length because it spans the 132 bp tandem direct repeats that differ in copy number between strains. The role of the 132 bp repeat region in the pathobiology of MD has been investigated thoroughly since the expansion of this region from 2 copies to over 20 occurs concomitantly with attenuation [46]. It has been reported that this expansion disrupts the 1.8 kb RNA transcript family that contains a putative fes/fps kinase-related transforming protein [47]; however, this expansion was proven insufficient to cause attenuation [48]. It was considered possible that this expansion affected the expression of proteins linked to the 1.8 kb RNA transcript (Fig 2A), but no evidence existed that the 132 bp direct repeats encoded a translated protein.

RNA-Seq results suggest MDV074, MDV075.4, and MDV075.7 at this locus are not expressed abundantly in epithelial skin cells (Fig S2, S3 Table), and the lack of matching peptides in MS/MS is consistent with this result. However, the region containing the MDV075.2 gene was abundantly expressed (3588 ± 461-fold coverage) in epithelial skin cells (S3 Table). The most likely reason for this depth of coverage is from non-spliced mRNAs of the highly abundant 14 kDa family of transcripts (discussed below) within an intron of which MDV075.2 is situated (Figs 2A, 6A). However, two unique peptides from the MDV075.2 protein sequence were identified (FLCLLPQGGGAR and RACS[+80]VTALAR) and provided support for the possibility that this variable-length ORF is translated into a protein product (Fig 2B). Annotated spectra and XIC for the first peptide are shown in S16A Fig. MS1 intensities for the peptide mass are low, but the XIC provides evidence that this ion is specific to infected samples. The matching b/y ions are moderate, although the strong y7 ion peak corresponding to an N-terminal proline agrees with the expected pattern from the known “proline effect” [49]. The XIC of the second peptide shows strong peaks in both infected and uninfected samples; this is likely a mis-assigned PSM (S16B Fig). Nevertheless, the data suggest the possibility that MDV075.2 within the 132 bp direct repeat is expressed, and the implications of the expansion of this region during attenuation are intriguing.

**Fig 6.**
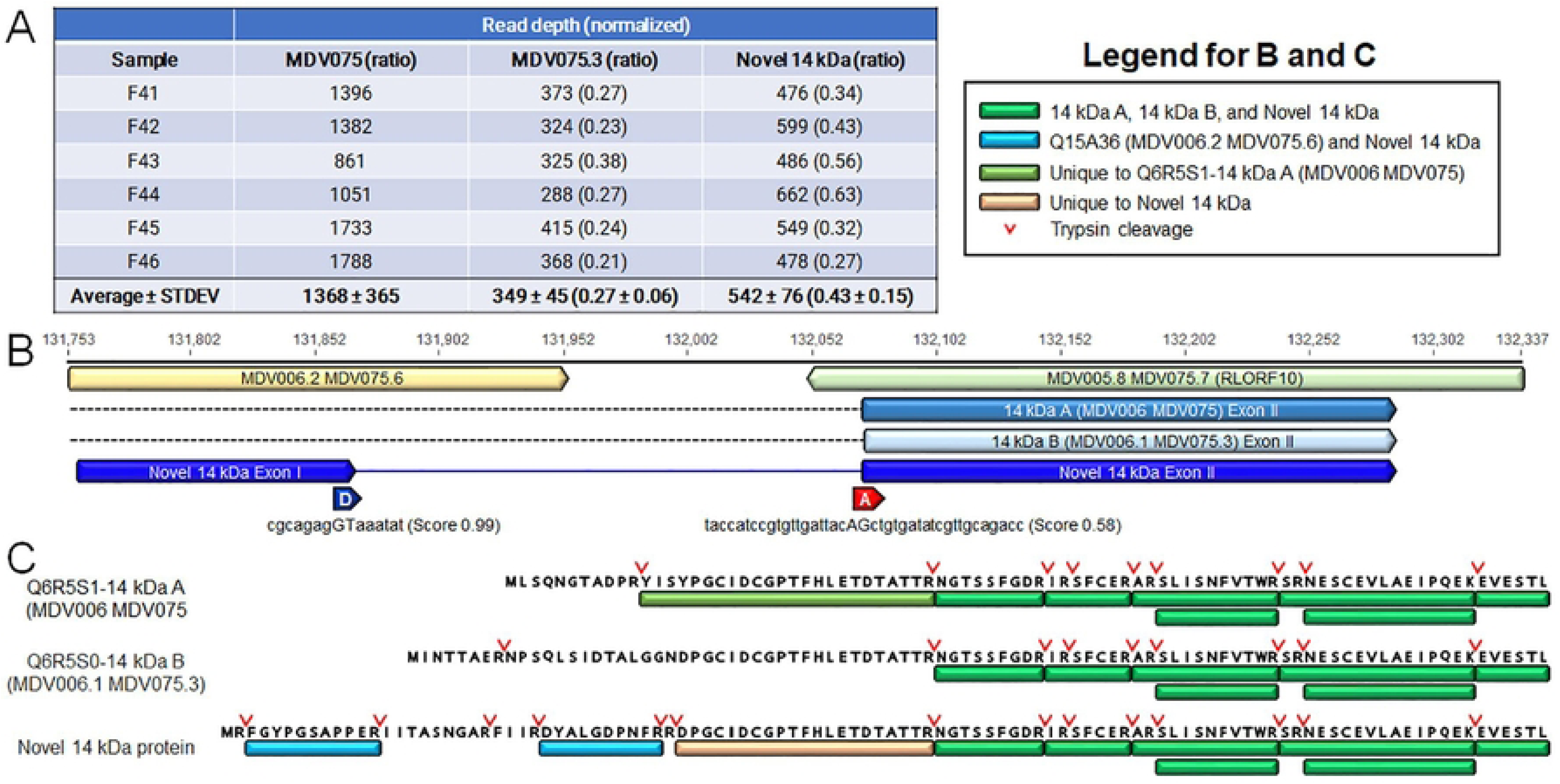
Expression of the 14 kDa family of genes, novel mRNA splicing, and validation of protein expression in epithelial skin cells. (A) Total reads for MDV075 (14 kDa A), MDV075.3 (14 kDa B), and the Novel 14 kDa transcripts for the six infected replicates with the average reads ± standard deviations in table form. Included is the ratio (in parentheses) of MDV075.3 (14 kDa B) and the novel 14 kDa transcripts compared to MDV075 (14 kDa A) ± standard deviations. (B) Schematic representation of MDV006.2 MDV075.6 and the 3’ end of the 1.8 kb transcript family encoding exons II of 14 kDa A, B, and Novel 14 kDa transcripts. Also included are the predicted donor (D) and acceptor (A) sites for the Novel 14 kDa intron using NNSPLICE 0.9 program [83] (C) Protein alignment of 14 kDa A, 14 kDa B, and Novel 14 kDa. Peptides unique to each protein are noted in the figure legend along with tryptic cleavage sites.

### Alternative splicing in the 14 kDa family of transcripts

The repeat long region of MDV has a complex arrangement of genes and expression patterns, including miRNAs [50, 51], internal ribosomal entry sites (IRES) [52], and circRNAs [53]. Of importance in the context of this report is the examination of mRNA splicing in the region spanning the 14 kDa family of nuclear proteins (Fig 2A, S2 Fig), whose transcripts are amongst the most highly expressed in epithelial skin cells (S3 Table). We detected abundant intron-spanning reads for both the known 14 kDa A and 14kDa B splice forms, in addition to a third splice variant with a novel exon I (Fig 6A). Exon I of this novel isoform is comprised of the 5’ half of the hypothetical ORF previously annotated as MDV075.6 (S2 Fig, Figs 2 & 6B). In epithelial skin cells, isoform A (MDV075) is the predominant RNA species, while the novel isoform is half as abundant, and isoform B is one-fourth as abundant as isoform A (Fig 6A).

Previous studies have shown that 14 kDa A and B were expressed at the protein level using polyclonal antibodies generated against each protein, but these studies could not differentiate 14 kDa A and B due to their shared protein sequences and lack of distinguishing peptide differences (39). We identified the intron-spanning peptide of the 14 kDa A isoform in the MS/MS dataset (Fig 3D) but not of the B isoform. Importantly, the proteogenomic scan identified the peptide spanning the splice junction of the novel 14 kDa isoform (DPGCIDCGPTFHLETDTATTR) shown in Figure 6C. This peptide is supported by rich MS2 spectra and infection-specific extracted ion chromatogram profiles (S17 Fig). It should be noted that the same peptide sequence is found spanning the B isoform splice junction, but in that protein it is not tryptic, as there is no Arg or Lys immediately upstream, and thus would not be detected in this assay (Fig 6C). Based on LFQ intensity, the A isoform-spanning peptide is ∼5× more abundant than the novel spanning peptide (4.5×10^6^ ± 9.1×10^5^ vs. 8.9×10^5^ ± 2.6×10^5^) (S3 Table). Both intron-spanning peptides were detected in all three infected replicates; neither was detected in any uninfected samples. Two additional peptides from all three infected replicates were identified from the previously annotated MDV075.6 ORF, now forming the 5’ exon of this novel splice product. Thus, the expression of the 14 kDa family of transcripts appears to be even more complicated as previously thought, but its expression in epithelial skins cells should be further investigated.

### Novel peptides identified by proteogenomic search

In addition to the novel translation start sites and spliced protein isoforms discovered by proteogenomic searching, several peptides were matched to the six-frame translation of the genome in previously unannotated ORFs (S7 Table). All these spectra passed the 1% PSM FDR threshold, but as with all novel peptides discussed herein, they require scrutiny. Of these, most were matched to single PSMs/replicates and have poor or inconclusive MS2 ion series and elution profiles. Two more (IS[+80]LNIR and TGN[+1]NISNNR) have strong elution peaks in both infected and uninfected replicates and are clearly misidentifications (S12B and C). However, the remaining three novel peptides are of interest.

The peptide EEFYEIYFEGCGSRSPTAR has an infection specific XIC but very sparse MS2 spectra (S18 Fig). However, it matches to the short protein sequence of the recently described SORF6 expressed transcript [17]. As in that publication, we also observed the splice junction associated with the SORF6 transcript in the RNA-Seq data (153 ± 30 spanning reads) and poly-A tailed reads mapping to the putative cleavage site downstream of a canonical polyadenylation signal. In light of a recent publication [26] in which the authors could not detect the translation of a tagged SORF6 coding sequence, the peptide evidence reported here, although not conclusive, beckons further exploration of the protein-coding potential of this ORF.

The two remaining novel peptides mapped to the 5’ ends of core genes on the same strand but out of frame. The peptide LEVDHAIVYR maps near the start of MDV060 (pUL47) within a short ORF having a start codon slightly upstream of the core gene (S19 Fig). There are two start codons upstream of the identified peptide, the furthest with a moderate Kozak consensus (AGT**ATG**C) and the second with a strong consensus (GGT**ATG**G). Starting from the upstream codon, a protein of 72 amino acids is predicted, with no known conserved functional motifs. This peptide has both a complete y-ion series and infection-specific XIC elution profiles (S19B Fig). Similarly, peptide FPAAPS[+80]PLPIAHAPVGLDSTR matches a small ORF overlapping the 5’ end of ICP4 (MDV084) (S20A Fig). This ORF has a *very* strong Kozak consensus (ACC**ATG**G) and codes for a putative 137 amino acid polypeptide with no known homology or functional motifs. This peptide also has reasonably strong support from the inspection of MS2 spectra and XIC (S20B Fig).

### Proteins notably missing in MS/MS

There were 84 proteins with at least one unique peptide detected. Of these, 4 are hypothetical proteins (MDV075.2, MDV075.6, MDV082, and MDV091.5). Our MS/MS analysis failed to detect peptides for 51 hypothetical and 17 annotated proteins (S3 Table). Of the ORFs with read coverage significantly above background but without peptides detected in MS/MS, several are of note. MDV015.5 (V57) lies at the 3’ end of the transcript containing MDV013 (UL1-gL), MDV014 (UL2), and MDV015 (UL3), all of which had high peptide coverage in this experiment (S1 Fig). We also detected a cluster of polyadenylated reads directly adjacent to a canonical polyadenylation signal downstream of MDV015.5, suggesting that it is likely transcribed as part of this gene locus. However, there are only two predicted tryptic peptides ≥ 6 aa in the UL15.5 protein sequence, making it reasonable to suppose that it was not detected by MS/MS due to the platform’s technical limitations. Similarly, MDV072.5 (UL56) lies on a transcript between MDV073 (pp38), MDV072 (LORF5), and MDV071 (CIRC), all of which have moderate to high peptide coverage (S1 Fig). This protein has only three predicted tryptic peptides ≥ 6 aa, from 27 to 58 aa long, again making it likely that it is missed due to technical limitations.

MDV056 (pUL43) also lies between two genes with high peptide coverage (pUL42 and pUL44- gC). In this case, unlike the short proteins above, it is predicted to generate 16 tryptic peptides (S21 Fig). However, it is a membrane protein of which 48% of the residues are predicted to lie within 11 transmembrane domains. It should be noted that Liu et al. [20] detected one peptide (MDSVNNSSLPPSYTTTGR) at the N-terminus of the protein in their study in cell culture. The overall hydrophobic nature of the protein may make it unamenable to detection with the methods used here. The same can be said for MDV032 (pUL20) (S22 Fig), although, unlike MDV056 (pUL43), the read depth for that gene in this experiment was barely above the baseline threshold (S3 Table). MDV075.8, located at the 3’ end of the 14 kDa protein family transcript, was another protein with high RNA-Seq levels but no peptide coverage. A deep cluster of polyadenylated reads shortly downstream of this ORF adds evidence that it is part of an abundant transcript. There are eight predicted detectable tryptic peptides in the protein sequence (data not shown), and it seems likely that it would have been detected if present at significant levels in the samples. Of note, this ORF contains a weak Kozak consensus (TGC**ATG**T), with conserved residues in neither the -3 nor +4 positions. The two remaining ORFs with high mRNA expression but no peptide coverage, MDV083, and MDV086, lie within the latency-associated transcript (LAT). Here, there is strong evidence of transcription in epithelial skin but no evidence of translation, in agreement with the accepted role of LAT as a non-coding transcript involved in transcriptional regulation [54].

Another protein not detected in our study was MDV035 (UL24), upstream of the well-represented UL25 (52% peptide coverage). Liu et al. [20] detected four unique peptides from this protein during cell culture replication, while Bertzbach et al. [17] did not. There are 22 predicted observable tryptic peptides in the protein (S23 Fig), and it is somewhat surprising that none were detected in epithelial skin cells despite relatively low mRNA levels (217 ± 33 fold-coverage). In other herpesvirus proteomics studies, this protein has also been difficult to detect [17, 20, 24, 25, 55]. As a rule, some proteins and peptides are inherently more difficult to detect using shotgun LC-MS/MS. Bell et al. failed to detect HSV-1 proteins UL11, UL20, UL43, and UL49.5 [24].

Loret *et al*. failed to detect HSV UL20, UL43, and UL49.5 (gN) in extracellular virions using shotgun proteomics, but using the more sensitive targeted multiple reaction monitoring technique were able to detect UL20 [56].

### Genes notably not expressed in epithelial skin cells

There were 38 “annotated” ORFs in this study with read depths below the threshold used as a measure of expression (S3 Table). Of these, 30 are annotated to encode “hypothetical proteins” that are nearly all < 200 aa in length (S24 Fig). Whether bona fide functional transcripts or not, our data suggest that they are most likely not expressed during fully productive viral replication in epithelial skin cells. It is possible some of these genes, particularly ORFs within the repeat regions, may be more robustly expressed in lymphocytes. However, Bertzbach *et al*. [17] also found many of these genes to be “not expressed” using an *in vitro* B cell infection model.

Of the remaining eight annotated ORFs with read depths below the threshold that have been characterized or at least described previously, the most notable are the two helicase-primase subunits, MDV017 (UL5) and MDV066 (UL52). Neither transcript is present above background levels, but both are clearly expressed in epithelial skin cells based on the multiple peptides detected in MS/MS (S2 & S3 Tables). However, their protein abundance is well below that of their neighboring genes, and it is likely that they are being expressed at low levels which would be more readily detectable under a different experimental approach. Similarly, MDV072 (LORF5) was detected by multiple MS/MS peptides and is also likely to be expressed in skin cells at low levels. For the remaining genes (MDV013.5, MDV057.8, MDV074, MDV075.7, MDV077) there was no evidence of translation, and they are unlikely to be expressed in epithelial skin cells. Notably, most are antisense to known expressed genes in the surrounding genomic context.

## Conclusions

This study provides a comprehensive analysis of the transcriptional and translational profile during fully productive skin-tropic herpesvirus replication in the host. To our knowledge, this is the first study in which both RNA-Seq and MS-based proteomics were employed in a natural herpesviral host model supporting fully productive virus replication. While we detected 114 viral ORFs with read depths above our threshold for expression, many of these are likely untranslated ORFs residing within transcribed loci. The 84 proteins (single peptide) or 79 proteins (2+ distinct peptides) detected with MS/MS are, therefore, likely a better representation of the protein-coding transcriptional landscape during fully productive replication.

We have demonstrated herein the application of a method to isolate virus-rich epithelial skin cell samples to maximize virus/host ratios and deeply integrate the proteome and transcriptome of productive infection. The demonstrated ability to reproducibly detect and quantify nearly all of the viral proteins expressed at this critical stage of infection, as well as obtain a high breadth of peptide coverage for many of them, opens up new possibilities to study viral protein functions in the natural host during fully productive herpesvirus replication whereby the effects of mutagenesis/perturbation at the protein and post-translational level can be directly studied. The viral enrichment strategy may further complement additional approaches, such as direct modified RNA sequencing and data-independent-acquisition MS/MS, to continue to push beyond simple gene expression and examine the finer aspects of virus/host molecular interactions.

## Materials and Methods

### Recombinant (r)MDV

The virus used here, vCHPKwt/10HA was recently reported [57], in which a 3×Flag and 2×HA epitopes were inserted in-frame of MDV UL13 (CHPK) and US10 at their C-termini, respectively, in addition to expressing pUL47eGFP [21].

### Ethics statement

All animal work was conducted according to national regulations. The animal care facilities and programs of UIUC meet all the requirements of the law (89 –544, 91–579, 94 –276) and NIH regulations on laboratory animals, comply with the Animal Welfare Act, PL 279, and are accredited by the Association for Assessment and Accreditation of Laboratory Animal Care (AAALAC). All experimental procedures were conducted in compliance with approved Institutional Animal Care and Use Committee protocols. Water and food were provided *ad libitum*.

### Animal experiments

Pure Columbian (PC) chickens were obtained from the UIUC Poultry Farm (Urbana, IL) and were from MD-vaccinated parents (Mab+). Twelve chicks were infected at three days of age with 2,000 PFU of cell-associated virus by intra-abdominal inoculation. Another fourteen age- matched, uninfected chicks were housed in a separate room.

To monitor the relative level of MDV in the chickens’ feathers during the infection, two flight feathers were plucked from each wing (4 total) starting at 14 days pi, fixed in 4% paraformaldehyde for 15 min, then washed twice with phosphate-buffered saline (PBS). Expression of pUL47eGFP was examined as previously described [58–60] using a Leica M205 FCA fluorescent stereomicroscope with a Leica DFC7000T digital color microscope camera (Leica Microsystems, Inc., Buffalo Grove, IL, USA). Chickens with heavily fluoresced feathers, along with age-matched uninfected birds, were euthanized to collect wing feathers for RNA and protein extractions [61]. All samples were collected between 21-35 days pi. Six chickens (replicates) of infected and uninfected groups were used for RNA sequencing, while three replicates of each group were used for LC/MS-MS.

### RNA extraction and RNA sequencing

The calamus of the feather tips collected were clipped with sterile scissors and dropped directly into 3.0 ml of RNA STAT-60 (Tel-Test, Inc., Friendswood, TX, USA), snap-frozen on dry ice, and stored at -80°C until all samples were collected. Samples were thawed at 37°C, mixed with a handheld homogenizer, and 1.0 ml transferred to Phasemaker Tubes (Invitrogen, Waltham, MA, USA) containing 200 µl of chloroform. The samples were vigorously mixed, incubated at room temperature for 3 min, then centrifuged (12,000 x *g* for 15 min at 4°C). Total RNA was precipitated with 500 µl isopropanol, washed with 75% ethanol, and dissolved in RNase-free water. The RNA quantity was determined using a Qubit RNA High Sensitivity Assay kit (Thermo Fisher, Suwanee, GA, USA), and its quality was determined using a Bioanalyzer 2100 (Agilent, Santa Clara, CA, USA). High-quality RNA samples with RIN values >7.0 were depleted of rRNAs using QIAseq FastSelect –rRNA HMR kit (Qiagen, Germantown, MD, USA) in combination with the KAPA stranded mRNA seq kit (Kapa Biosystems, Wilmington, MA, USA). Ribo-depleted RNA was suspended in the Fragment/Prime/Elute mix and fragmented at 94°C for eight min. Using the same KAPA kit, cDNAs were generated using random hexamer priming, end-repaired, and indexed with individual adaptors. Libraries were quantified using a Qubit fluorometer and analyzed on a Bioanalyzer 2100 to determine the size distribution of the library. Pooling cDNAs with fragments (200–300 bp) was done using qPCR concentrations. The quality of the final pool was determined using Qubit, fragment analyzer, and qPCR. RNA libraries were prepared for sequencing on Illumina NextSeq 500 instrument using Illumina’s dilute and denature protocol. Pooled libraries were diluted to 2nM, then denatured using NaOH. The denatured libraries were further diluted to 2.2pM, and PhiX was added to 1% of the library volume. Data were demultiplexed and trimmed of adapter sequences, and barcoded sequences were uploaded onto the BaseSpace Sequencing Hub.

### RNA-Seq data analysis

#### Visualization of RNA sequencing and proteomics in IGV

Genome tracks and additional data tracks generated from this study were visualized using IGV [62]. Online visualization of the data tracks was build using igv.js [63] and is accessible at https://igv.base2.bio/AAG3-9Fja-99a2-2asZ/.

#### Preprocessing and read mapping

Raw paired RNA-Seq reads were preprocessed using Trim Galore v. 0.6.6 (https://www.bioinformatics.babraham.ac.uk/projects/trim_galore/) in two-color mode, minimum quality 8, minimum trimmed length 40, automatic adapter detection. Reads were mapped against the combined host (bGalGal1.mat.broiler.GRCg7b) and RB-1B (modified from MT272733 based on epitope tags incorporated) genomes using the splice-aware mapper HISAT2 v. 2.2.1 [64], maximum intron length = 50000, strandedness = RF. The RB-1B reference used throughout was trimmed to remove redundant terminal repeat sequences, except for short regions surrounding the TRL/UL and US/TRS junctions which were included to retain putative junction-spanning genes. Strandedness efficiency was calculated using the infer_experiment.py script from RSeqC v. 4.0.0 [65]. The resulting alignment files were filtered for reads mapping to the viral genome and split into four strand-specific subsets (forward read/ forward strand, forward read/reverse strand, reverse read/forward strand, and reverse read/reverse strand) using SAMtools v. 1.14 [66], filtering on the SAM flags 0×10, 0×40, and 0×80. For all further steps, only the six infected replicates were used (read counts from the uninfected replicates mapping to the MDV genome were zero or near-zero - see S4 Table). Read-spanning intron junctions were extracted from the BAM alignments using the RegTools [67] “junctions extract” command (-a 15 -m 20 -M 50000 -s 1). Only junctions supported by ten or more reads per replicate were used for further analysis and visualization.

#### Gene read coverage and background calculation

Because a high degree of intergenic and/or non-strand-specific mapping to the viral genome was detected in preliminary analysis (see results for further details), it was decided to use background-subtracted median read depth per gene as a metric for evaluating gene transcriptional status. To this end, strand-specific per-base read depth in bedgraph format was calculated using the BEDTools v. 2.30.0 tool genomecov [68]. Strand-specific per-sample median read depths for each annotated MDV gene were then calculated using the BEDTools map command. Inter-sample normalization of gene read depths was performed using median centering, after which overall means, and standard deviations were calculated for each gene. Calculation of the background/non-specific read depth was performed as follows. First, because the untranslated regions of the MDV gene models are not well-defined, gene intervals were estimated by adding 600 bp upstream and 100 bp downstream of the annotated coding sequence coordinates using the BEDTools “slop” command. This file was used to mask the full genome coverage bedgraph file using the BEDTools “subtract” command, resulting in an array of per-base intergenic/antisense read depths. Because the distribution of these values was assumed to be a mixture of at least two groups (a large group of true intergenic/antisense positions and a smaller group of actually transcribed positions), the “normalmixEM” method from the R mixtools package v. 1.2.0 [69] was used to generate a preliminary parameter estimate of the true intergenic read depth distribution, assuming a Gaussian distribution. These estimates were further adjusted manually by visualization in R to the final values of mean 50×, standard deviation 33× (S9 Fig). A threshold read depth of mean plus two standard deviations (116×) was used subsequently to determine genes which were expressed above background levels with ∼98% confidence (single-tailed).

### Protein extraction, proteomics, and phosphopeptide analyses

Feathers from MDV-infected and age-matched uninfected birds were plucked, placed into ice-cold PBS, and epithelial skin scrapings were provided to the University of Illinois Protein Sciences Facility as frozen samples. They were subsequently lysed in a buffer containing 6 M guanidine HCl, ten mM tris(2-carboxyethyl)phosphine HCL, 40 mM 2-chloroacetamide, and 0.1% sodium deoxycholate and then boiled to promote reduction and alkylation of disulfide bonds, as previously described [70]. The samples were cleared of debris by centrifugation and subjected to chloroform-methanol precipitation to remove lipids and other impurities; the resulting protein pellets were dissolved in 100 mM triethylammonium bicarbonate with sonication. Protein amounts were determined by BCA assay (Pierce, Rockford, IL) before sequential proteolytic digestion by LysC (1:100 w/w enzyme: substrate; Wako Chemicals, Richmond, VA) for 4 h at 30°C and trypsin (1:50 w/w; Pierce) overnight at 37°C. Peptide samples were desalted using Sep-Pak C18 columns (Waters, Milford, MA) and dried in a vacuum centrifuge. For phosphorylation analysis, phosphopeptides were enriched by iron- immobilized metal ion affinity chromatography (Fe-IMAC) in a microtip format before being desalted once more using StageTips [71].

Peptide digests were analyzed using a Thermo UltiMate 3000 UHPLC system coupled to a high resolution Thermo Q Exactive HF-X mass spectrometer. Peptides were separated by reversed-phase chromatography using a 25 cm Acclaim PepMap 100 C18 column maintained at 50°C with mobile phases of 0.1% formic acid (A) and 0.1% formic acid in 80% acetonitrile (B). A two-step linear gradient from 5% B to 35% B over the course of 110 min and 35% B to 50% B over 10 min was employed for peptide separation, followed by additional steps for column washing and equilibration. The MS was operated in a data-dependent manner in which precursor scans from 350 to 1500 m/z (120,000 resolution) were followed by higher-energy collisional dissociation (HCD) of the 15 most abundant ions. MS2 scans were acquired at a resolution of 15,000 with a precursor isolation window of 1.2 m/z and a dynamic exclusion window of 60 s.

The raw LC-MS/MS data was analyzed against the Uniprot GaHV2 database (taxon 10390; 1300 sequences) using the Byonic peptide search algorithm (Protein Metrics) integrated into Proteome Discoverer 2.4 (Thermo Scientific). Optimal main search settings were initially determined with Byonic Preview (Protein Metrics) and included a peptide precursor mass tolerance of 8 ppm with fragment mass tolerance of 20 ppm. Tryptic digestion was specified with a maximum of 2 missed cleavages. Variable modifications included oxidation/dioxidation of methionine, acetylation of protein N-termini, deamidation of asparagine, conversion of peptide N-terminal glutamic acid/glutamine to pyroglutamate, and phosphorylation of serine, threonine, and tyrosine. A static modification to account for cysteine carbamidomethylation was also added to the search. PSM false discovery rates were estimated by Byonic using a target/decoy approach.

### Additional proteogenomic analysis

To search for potential novel expressed reading frames and proteoforms, three additional MS/MS search databases were generated from the rRB-1B genome sequence using in-house software. A database of tryptic peptides spanning all putative transcript splice sites identified from RNA-Seq was generated, adding an ambiguous residue (X) at each end to prevent the search engine from assuming a protein terminus. A second database containing a full six-frame translation of the genome, split at stop codons, was generated, again adding an ambiguous base at the N-terminus to prevent identification as a protein terminus. A third database was generated containing possible alternative N-terminal peptides based on potential alternative translation initiation sites (TIS) as follows. For each annotated gene model, all in-frame moderate Kozak consensus sequences (A|G at -3 position, ATG|CTG|GTG|ACG|ATA|TTG|ATT at +1-3, G at +4) were identified between the annotated TIS and the first in-frame upstream stop codon, and a putative tryptic peptide was added to the database for each one after replacing the first amino acid (for non-canonical start codons) with methionine. A similar scan was performed for alternative downstream TIS, limited to a maximum of four.

These three additional databases were combined with databases of the annotated RB-1B proteins, the annotated host proteins from chicken genome assembly bGalGal1.mat.broiler.GRCg7b, and the cRAP database of common contaminant proteins (https://www.thegpm.org/crap/), along with reversed decoy sequences of each entry. Raw spectra were searched against this database using Comet v. 2019.01 rev. 5 [72], MS-GF+ v. 2022.01.07 [73], and Byonic as described above.

Search parameters included a precursor mass tolerance of 7 ppm; high-resolution MS2 mass tolerance (MS-GF+ InstrumentID=1, Comet fragment_bin_tol=0.02 + fragment_bin_offset=0.0); fully-tryptic termini; maximum two missed cleavages; fixed Cys carbamidomethylation; variable S/T/Y phosphorylation, Met oxidation, N/Q deadmidation, N-terminal protein acetylation, and N-terminal methionine excision. Raw spectral hits were post-processed using Percolator v. 3.05 [74]to assign q-values at the spectrum, peptide, and protein levels for use in false discovery rate (FDR) filtering. Comet and Percolator were run within the Crux toolkit v. 4.1 [75]. Visualization of identified peptides was performed in IGV [62]. All search databases and Crux and MS-GF+ configuration files are available upon request.

Peptide intensity calculation was performed using FlashLFQ v. 1.2.4 [76] with match-between-run (MBR) enabled, inter-sample normalization, and requiring MS2 ID in condition for MBR. Peptide intensities were used to calculate protein iBAQ values by dividing summed peptide intensities for each protein by the number of theoretical fully tryptic peptides length 6-40 in the protein. Intensities for peptides shared between proteins were divided evenly between proteins. Relative iBAQ (riBAQ) was calculated within each replicate as the protein iBAQ divided by the sum of iBAQ values for the replicate, considering only viral proteins.

### Statistical analysis

Statistical analysis of the RNA-Seq and proteomics data was performed using the R software package v. 4.1.3 [77]. Analysis of proteomics data within R was partially performed using the MSnbase package v. 2.20.4 [78]. Visualizations of annotated MS2 spectra and combined extracted ion chromatograms (XICs) were created using ms-perl (https://metacpan.org/pod/MS) and R. For the purpose of six-replicate XICs, the raw data were aligned across retention times using the MapAlignerPoseClustering tool from OpenMS [79], with superimposer:mz_pair_max_distance=0.05 and pairfinder:distance_MZ:max_difference = 7 ppm.

## Acknowledgements

We thank the Georgia Genomics and Bioinformatics Core, which provided the Illumina: Ribo-depleted RNA library preparation and NextSeq500 2×75bp sequencing service and the University of Illinois at Urbana-Champaign Protein Sciences core for their guidance in LC-MS/MS services. This report was supported by Agriculture and Food Research Initiative Competitive Grant nos. 2013-67015-26787, 2016-67015-26777, and 2020-67015-21399 from the USDA National Institute of Food and Agriculture, and USDA-ARS NACA agreements nos. 58- 6040-8-037 and 58-6040-0-015.

## Supporting information captions

**S1 Fig. The transcriptome and proteome of the unique long (UL) region of MDV in epithelial skin cells.** Visualization of the introns, genes, log2 read depth, and peptides detected on the forward (blue) and complementary (red) strands within the UL region plus 500 bp flanking sequences of MDV during fully productive replication.

**S2 Fig. The transcriptome and proteome of the repeat long (RL) region of MDV in epithelial skin cells.** Visualization of the introns, genes, log2 read depth, and peptides detected on the forward (blue) and complementary (red) strands within the RL region plus 500 bp flanking sequences of MDV during fully productive replication.

**S3 Fig. The transcriptome and proteome of the repeat short (RS) region of MDV in epithelial skin cells.** Visualization of the introns, genes, log2 read depth, and peptides detected on the forward (blue) and complementary (red) strands within the RS region plus 500 bp flanking sequences of MDV during fully productive replication.

**S4 Fig. The transcriptome and proteome of the unique short (US) region of MDV in epithelial skin cells.** Visualization of the introns, genes, log2 read depth, and peptides detected on the forward (blue) and complementary (red) strands within the US region plus 500 bp flanking sequences of MDV during fully productive replication.

**S5 Fig. Unique peptide counts and breadth of coverage for detected viral proteins in infected epithelial skin cells.**

**S6 Fig. Evidence of expression of MDV023 (pUL11) in infected cells based on a single peptide.** (A) Protein sequence of MDV023 (pUL11) comparing the reference sequence (G9CUB8) and the RB-1B strain used in this study, plus the predicted tryptic cleavage sites. (B) Elution peaks of XIC showing unique spectra in replicates of infected samples relative to uninfected samples.

**S7 Fig. Evidence for expression of MDV064 (pUL49.5-gN) in infected cells based on a single peptide.** (A) Protein sequence of MDV064 (pUL49.5-gN) comparing the reference sequence (QM77MR4) and the RB-1B strain used in this study, plus the predicted tryptic cleavage sites, predicted signal peptide and transmembrane regions. (B) Elution peaks of XIC showing unique spectra in replicates of infected samples relative to uninfected samples.

**S8 Fig. Evidence against expression of Tryptic map and XTC spectra of single peptides for MDV096 (Meq), MDV094 (SORF4), and MDV091.5.** The XTC spectra of single unique peptides detected for MDV096 (A), MDV094 (B), and MDV0915 (C) showing little confidence in their specificity.

**S9 Fig. Baseline read depth distribution calculated for intergenic/antisense regions.**

**S10 Fig. Correlation between RNA and protein levels.**

**S11 Fig. Rich y/b ion series and infection-specific elution profiles of exon spanning peptides for MDV078-vCXCL13 (A), MDV057.1-gC104 (B), and MDV075-14 kDa A (C).**

**S12 Fig. Poor y/b ion series and infection-specific elution profiles of peptides.**

**S13 Fig. Alternative TIS for MDV055 (pUL42).** (A) 5’ end of MDV055 showing N-terminal peptides for both the annotated and alternative TIS identified by peptides. (B & C) The annotated (B) and alternative (C) y/b ion series and elution profiles are shown.

**S14 Fig. Alternative TIS for MDV070 (pUL55).** (A) 5’ end of MDV070 showing N-terminal peptides for both the annotated and alternative TIS identified by peptides. (B & C) The annotated (B) and alternative (C) y/b ion series and elution profiles are shown.

**S15 Fig. Alternative splicing of MDV008/MDV073.** (A) MDV008 and MDV073 overlap the junction between the UL and RL regions creating alternative proteins including previously identified pp38 and pp24, and pp38A and pp38B created through alternative splicing. A novel splice variant termed Novel pp38C is expressed in epithelial skin cells. Donor (D) and acceptor (A) locations are shown. (B) MUSCLE alignment of pp38A, pp38B, and Novel pp38C with trypsin cleavage sites. (C) MUSCLE alignment of pp38, pp24, pp38A, pp38B, and Novel pp38C and peptides detected in epithelial skin cells. Some peptides are unique to specific proteins.

**S16 Fig. Y/b ion series and elution profiles for peptides spanning MDV075.2.**

**S17 Fig. Rich y/b ion series and infection-specific elution profiles for novel 14 kDa isoform.**

**S18 Fig. Tryptic map and elution profiles for SORF6.** (A) Exon I and II of SORF6, location of tryptic cleavage sites, and peptides identified in infected samples. (B &C) Elution profiles for SORF6 peptide.

**S19 Fig. Novel microORF within MDV060.** (A) 5’ region of MDV060 with the coding sequence, TIS for pUL47, and peptides mapping to annotated pUL47. A novel peptide was detected using 6-frame translation identified a novel microORF. Two potential TIS for the novel microORF are shown. (B) Elution profiles for the 6-frame peptide in all three infected samples.

**S20 Fig. Novel microORF within MDV084 (ICP4).** (A) 5’ region of MDV084 with the coding sequence, TIS for ICP4, and peptides mapping to annotated ICP4 in green. A novel peptide (blue) was detected using 6-frame translation identified a novel microORF (orange). B) Elution profiles for the 6-frame peptide in all three infected samples.

**S21 Fig. Tryptic map for MDV056 (pUL43).** Protein sequence of MDV056 (pUL43) comparing the reference sequence (Q9E6M9) and the RB-1B strain used in this study, plus the predicted tryptic cleavage sites. Transmembrane regions and the unique peptide identified in Liu et al. [20] are shown.

**S22 Fig. Tryptic map for MDV032 (pUL20).** Protein sequence of MDV032 (pUL20) comparing the reference sequence (Q77MS4) and the RB-1B strain used in this study. The predicted tryptic cleavage sites and transmembrane regions are shown.

**S23 Fig. Tryptic map for MDV035 (pUL24).** Protein sequence of MDV035 (pUL24) comparing the reference sequence (Q9E6P4) and the RB-1B strain used in this study. The predicted tryptic cleavage sites and unique peptides identified in Liu et al. [20] are shown.

**S24 Fig. Predicted protein sizes for annotated MDV genes.**

**S1 Table. Total viral proteins detected in unenriched and phospho-enriched protein extracts of three uninfected and MDV-infected samples.**

**S2 Table. The peptide-spectrum matches for viral peptides identified, samples and engines identified in, including annotated, alternative starts, and 6-frame translation.**

**S3 Table. Summary of viral RNA and proteins in epithelial skin cells.** Worksheets show viral genes detected at both RNA and protein level, RNA only, and not detected in both RNA seq and LC-MS/MS.

**S4 Table. Summary of viral RNA sequencing data in epithelial skin cells. S5 Table. N-terminal peptides detected in epithelial skin cells.**

**S6 Table. LFQ intensities for peptides detected in epithelial skin cells. S7 Table. Peptides detected using six-frame proteogenomic searching.**

## References

1. Roizman B, Knipe DM, Whitley RJ. Herpes simplex viruses. In: Knipe DM, Howley PM, editors. Fields Virology. 2. 5th ed. Philadelphia, PA: Lippincott Williams & Wilkins; 2007. p. 2501–601.

2. Okura T, Taneno A, Oishi E. Cell-to-Cell Transmission of Turkey Herpesvirus in Chicken Embryo Cells via Tunneling Nanotubes. Avian Dis. 2021;65(3):335–9. doi: 10.1637/aviandiseases-D-21-00022.

3. Davison AJ. Evolution of the herpesviruses. Vet Microbiol. 2002;86(1-2):69–88.

4. Davison AJ, Dargan DJ, Stow ND. Fundamental and accessory systems in herpesviruses. Antiviral Res. 2002;56(1):1–11. Epub 2002/09/27.

5. Couteaudier M, Courvoisier K, Trapp-Fragnet L, Denesvre C, Vautherot JF. Keratinocytes derived from chicken embryonic stem cells support Marek’s disease virus infection: a highly differentiated cell model to study viral replication and morphogenesis. Virol J. 2016;13(1):7. doi: 10.1186/s12985-015-0458-2.

6. Wen L, Zhang A, Li Y, Lai H, Li H, Luo Q, et al. Suspension culture of Marek’s disease virus and evaluation of its immunological effects. Avian pathology : journal of the WVPA. 2019;48(3):183–90. doi: 10.1080/03079457.2018.1556385.

7. Li X, Schat KA. Quail cell lines supporting replication of Marek’s disease virus serotype 1 and 2 and herpesvirus of turkeys. Avian Dis. 2004;48(4):803–12. doi: 10.1637/7182-032604R.

8. Geerligs H, Spijkers I, Rodenberg J. Efficacy and safety of cell-associated vaccines against Marek’s disease virus grown in QT35 cells or JBJ-1 cells. Avian Dis. 2013;57(2 Suppl):448-53. doi: 10.1637/10344-090312-Reg.1.

9. Geerligs H, Quanz S, Suurland B, Spijkers IE, Rodenberg J, Davelaar FG, et al. Efficacy and safety of cell associated vaccines against Marek’s disease virus grown in a continuous cell line from chickens. Vaccine. 2008;26(44):5595–600. doi: 10.1016/j.vaccine.2008.07.080.

10. Abujoub A, Coussens PM. Development of a sustainable chick cell line infected with Marek’s disease virus. Virology. 1995;214(2):541–9. doi: 10.1006/viro.1995.0065.

11. Calnek BW, Adldinger HK, Kahn DE. Feather follicle epithelium: a source of enveloped and infectious cell-free herpesvirus from Marek’s disease. Avian Dis. 1970;14(2):219–33. doi: 10.2307/1588466.

12. Jarosinski KW, Margulis NG, Kamil JP, Spatz SJ, Nair VK, Osterrieder N. Horizontal transmission of Marek’s disease virus requires US2, the UL13 protein kinase, and gC. J Virol. 2007;81(19):10575–87. Epub 20070718. doi: 10.1128/JVI.01065-07.

13. Tulman ER, Afonso CL, Lu Z, Zsak L, Rock DL, Kutish GF. The genome of a very virulent Marek’s disease virus. J Virol. 2000;74(17):7980–8.

14. Lee LF, Wu P, Sui D, Ren D, Kamil J, Kung HJ, et al. The complete unique long sequence and the overall genomic organization of the GA strain of Marek’s disease virus. Proc Natl Acad Sci U S A. 2000;97(11):6091–6. Epub 2000/05/24. doi: 10.1073/pnas.97.11.6091.

15. Osterrieder N, Vautherot JF. The genome content of Marek’s disease-like viruses. In: Davison TF, Nair VK, editors. Marek’s disease. Biology of Animal Infection. London: Elsevier; 2004. p. 17–31.

16. Sadigh Y, Tahiri-Alaoui A, Spatz S, Nair V, Ribeca P. Pervasive Differential Splicing in Marek’s Disease Virus can Discriminate CVI-988 Vaccine Strain from RB-1B Very Virulent Strain in Chicken Embryonic Fibroblasts. Viruses. 2020;12(3). doi: 10.3390/v12030329.

17. Bertzbach LD, Pfaff F, Pauker VI, Kheimar AM, Hoper D, Hartle S, et al. The Transcriptional Landscape of Marek’s Disease Virus in Primary Chicken B Cells Reveals Novel Splice Variants and Genes. Viruses. 2019;11(3). doi: 10.3390/v11030264.

18. Sunkaraa L, Ahmad SM, Heidari M. RNA-seq analysis of viral gene expression in the skin of Marek’s disease virus infected chickens. Vet Immunol Immunopathol. 2019;213:109882. doi: 10.1016/j.vetimm.2019.109882.

19. Buza JJ, Burgess SC. Modeling the proteome of a Marek’s disease transformed cell line: a natural animal model for CD30 overexpressing lymphomas. Proteomics. 2007;7(8):1316–26. doi: 10.1002/pmic.200600946.

20. Liu HC, Soderblom EJ, Goshe MB. A mass spectrometry-based proteomic approach to study Marek’s Disease Virus gene expression. J Virol Methods. 2006;135(1):66–75. doi: 10.1016/j.jviromet.2006.02.001.

21. Jarosinski KW, Arndt S, Kaufer BB, Osterrieder N. Fluorescently tagged pUL47 of Marek’s disease virus reveals differential tissue expression of the tegument protein in vivo. J Virol. 2012;86(5):2428–36. Epub 20111221. doi: 10.1128/JVI.06719-11.

22. Jarosinski KW, Osterrieder N. Marek’s disease virus expresses multiple UL44 (gC) variants through mRNA splicing that are all required for efficient horizontal transmission. J Virol. 2012;86(15):7896–906. Epub 20120516. doi: 10.1128/JVI.00908-12.

23. Jarosinski KW. Marek’s disease virus late protein expression in feather follicle epithelial cells as early as 8 days postinfection. Avian Dis. 2012;56(4):725–31. doi: 10.1637/10252-052212-Reg.1.

24. Bell C, Desjardins M, Thibault P, Radtke K. Proteomics analysis of herpes simplex virus type 1-infected cells reveals dynamic changes of viral protein expression, ubiquitylation, and phosphorylation. J Proteome Res. 2013;12(4):1820–9. Epub 20130304. doi: 10.1021/pr301157j.

25. Ouwendijk WJD, Dekker LJM, van den Ham HJ, Lenac Rovis T, Haefner ES, Jonjic S, et al. Analysis of Virus and Host Proteomes During Productive HSV-1 and VZV Infection in Human Epithelial Cells. Front Microbiol. 2020;11:1179. Epub 20200529. doi: 10.3389/fmicb.2020.01179.

26. You Y, Conradie AM, Kheimar A, Bertzbach LD, Kaufer BB. The Marek’s Disease Virus Unique Gene MDV082 Is Dispensable for Virus Replication but Contributes to a Rapid Disease Onset. J Virol. 2021;95(15):e0013121. Epub 20210712. doi: 10.1128/JVI.00131-21.

27. Li X, Jarosinski KW, Schat KA. Expression of Marek’s disease virus phosphorylated polypeptide pp38 produces splice variants and enhances metabolic activity. Vet Microbiol. 2006;117(2-4):154–68. Epub 20060727. doi: 10.1016/j.vetmic.2006.06.019.

28. Jarosinski KW, Schat KA. Multiple alternative splicing to exons II and III of viral interleukin-8 (vIL-8) in the Marek’s disease virus genome: the importance of vIL-8 exon I. Virus Genes. 2007;34(1):9–22. Epub 20060822. doi: 10.1007/s11262-006-0004-9.

29. Parcells MS, Lin SF, Dienglewicz RL, Majerciak V, Robinson DR, Chen HC, et al. Marek’s disease virus (MDV) encodes an interleukin-8 homolog (vIL-8): characterization of the vIL-8 protein and a vIL-8 deletion mutant MDV. J Virol. 2001;75(11):5159–73. doi: 10.1128/JVI.75.11.5159-5173.2001.

30. Peng Q, Shirazi Y. Isolation and characterization of Marek’s disease virus (MDV) cDNAs from a MDV-transformed lymphoblastoid cell line: identification of an open reading frame antisense to the MDV Eco-Q protein (Meq). Virology. 1996;221(2):368–74. doi: 10.1006/viro.1996.0388.

31. Cantello JL, Parcells MS, Anderson AS, Morgan RW. Marek’s disease virus latency-associated transcripts belong to a family of spliced RNAs that are antisense to the ICP4 homolog gene. J Virol. 1997;71(2):1353–61. doi: 10.1128/JVI.71.2.1353-1361.1997.

32. Peng Q, Zeng M, Bhuiyan ZA, Ubukata E, Tanaka A, Nonoyama M, et al. Isolation and characterization of Marek’s disease virus (MDV) cDNAs mapping to the BamHI-I2, BamHI-Q2, and BamHI-L fragments of the MDV genome from lymphoblastoid cells transformed and persistently infected with MDV. Virology. 1995;213(2):590–9. doi: 10.1006/viro.1995.0031.

33. Hong Y, Coussens PM. Identification of an immediate-early gene in the Marek’s disease virus long internal repeat region which encodes a unique 14-kilodalton polypeptide. J Virol. 1994;68(6):3593–603.

34. Grzegorski SJ, Chiari EF, Robbins A, Kish PE, Kahana A. Natural variability of Kozak sequences correlates with function in a zebrafish model. PLoS One. 2014;9(9):e108475. Epub 20140923. doi: 10.1371/journal.pone.0108475.

35. Wingfield PT. N-Terminal Methionine Processing. Curr Protoc Protein Sci. 2017;88:6 14 1-6 3. Epub 20170403. doi: 10.1002/cpps.29.

36. Sedlackova L, Perkins KD, Lengyel J, Strain AK, van Santen VL, Rice SA. Herpes simplex virus type 1 ICP27 regulates expression of a variant, secreted form of glycoprotein C by an intron retention mechanism. J Virol. 2008;82(15):7443–55. doi: 10.1128/JVI.00388-08.

37. Schat KA, Piepenbrink MS, Buckles EL, Schukken YH, Jarosinski KW. Importance of differential expression of Marek’s disease virus gene pp38 for the pathogenesis of Marek’s disease. Avian Dis. 2013;57(2 Suppl):503–8. doi: 10.1637/10414-100612-Reg.1.

38. Deckman IC, Hagen M, McCann PJ, 3rd. Herpes simplex virus type 1 protease expressed in Escherichia coli exhibits autoprocessing and specific cleavage of the ICP35 assembly protein. J Virol. 1992;66(12):7362–7. doi: 10.1128/JVI.66.12.7362-7367.1992.

39. Liu F, Roizman B. Characterization of the protease and other products of amino-terminus-proximal cleavage of the herpes simplex virus 1 UL26 protein. J Virol. 1993;67(3):1300–9. doi: 10.1128/JVI.67.3.1300-1309.1993.

40. Weinheimer SP, McCann PJ, 3rd, O’Boyle DR, 2nd, Stevens JT, Boyd BA, Drier DA, et al. Autoproteolysis of herpes simplex virus type 1 protease releases an active catalytic domain found in intermediate capsid particles. J Virol. 1993;67(10):5813–22. doi: 10.1128/JVI.67.10.5813-5822.1993.

41. DiIanni CL, Drier DA, Deckman IC, McCann PJ, 3rd, Liu F, Roizman B, et al. Identification of the herpes simplex virus-1 protease cleavage sites by direct sequence analysis of autoproteolytic cleavage products. The Journal of biological chemistry. 1993;268(3):2048–51.

42. Gao M, Matusick-Kumar L, Hurlburt W, DiTusa SF, Newcomb WW, Brown JC, et al. The protease of herpes simplex virus type 1 is essential for functional capsid formation and viral growth. J Virol. 1994;68(6):3702–12. doi: 10.1128/JVI.68.6.3702-3712.1994.

43. Davison MD, Rixon FJ, Davison AJ. Identification of genes encoding two capsid proteins (VP24 and VP26) of herpes simplex virus type 1. J Gen Virol. 1992;73 (Pt 10):2709–13. doi: 10.1099/0022-1317-73-10-2709.

44. Reese MG. Application of a time-delay neural network to promoter annotation in the Drosophila melanogaster genome. Comput Chem. 2001;26(1):51–6. doi: 10.1016/s0097-8485(01)00099-7.

45. Liu FY, Roizman B. The herpes simplex virus 1 gene encoding a protease also contains within its coding domain the gene encoding the more abundant substrate. J Virol. 1991;65(10):5149–56. doi: 10.1128/JVI.65.10.5149-5156.1991.

46. Maotani K, Kanamori A, Ikuta K, Ueda S, Kato S, Hirai K. Amplification of a tandem direct repeat within inverted repeats of Marek’s disease virus DNA during serial in vitro passage. J Virol. 1986;58(2):657–60.

47. Peng F, Bradley G, Tanaka A, Lancz G, Nonoyama M. Isolation and characterization of cDNAs from BamHI-H gene family RNAs associated with the tumorigenicity of Marek’s disease virus. J Virol. 1992;66(12):7389–96. doi: 10.1128/JVI.66.12.7389-7396.1992.

48. Silva RF, Reddy SM, Lupiani B. Expansion of a unique region in the Marek’s disease virus genome occurs concomitantly with attenuation but is not sufficient to cause attenuation. J Virol. 2004;78(2):733–40.

49. Breci LA, Tabb DL, Yates JR, 3rd, Wysocki VH. Cleavage N-terminal to proline: analysis of a database of peptide tandem mass spectra. Analytical chemistry. 2003;75(9):1963–71. doi: 10.1021/ac026359i.

50. Burnside J, Bernberg E, Anderson A, Lu C, Meyers BC, Green PJ, et al. Marek’s disease virus encodes MicroRNAs that map to meq and the latency-associated transcript. J Virol. 2006;80(17):8778–86. doi: 10.1128/JVI.00831-06.

51. Yao Y, Zhao Y, Xu H, Smith LP, Lawrie CH, Sewer A, et al. Marek’s disease virus type 2 (MDV-2)-encoded microRNAs show no sequence conservation with those encoded by MDV- 1. J Virol. 2007;81(13):7164–70. doi: 10.1128/JVI.00112-07.

52. Tahiri-Alaoui A, Matsuda D, Xu H, Panagiotis P, Burman L, Lambeth LS, et al. The 5’ leader of the mRNA encoding the marek’s disease virus serotype 1 pp14 protein contains an intronic internal ribosome entry site with allosteric properties. J Virol. 2009;83(24):12769–78. doi: 10.1128/JVI.01010-09.

53. Chasseur AS, Trozzi G, Istasse C, Petit A, Rasschaert P, Denesvre C, et al. Marek’s Disease Virus Virulence Genes Encode Circular RNAs. J Virol. 2022:e0032122. Epub 20220412. doi: 10.1128/jvi.00321-22.

54. Kennedy PG, Rovnak J, Badani H, Cohrs RJ. A comparison of herpes simplex virus type 1 and varicella-zoster virus latency and reactivation. J Gen Virol. 2015;96(Pt 7):1581–602. Epub 20150320. doi: 10.1099/vir.0.000128.

55. Johannsen E, Luftig M, Chase MR, Weicksel S, Cahir-McFarland E, Illanes D, et al. Proteins of purified Epstein-Barr virus. Proc Natl Acad Sci U S A. 2004;101(46):16286–91. Epub 20041108. doi: 10.1073/pnas.0407320101.

56. Loret S, Guay G, Lippe R. Comprehensive characterization of extracellular herpes simplex virus type 1 virions. J Virol. 2008;82(17):8605–18. doi: 10.1128/JVI.00904-08.

57. Ponnuraj N, Akbar H, Arrington JV, Spatz SJ, Nagarajan B, Desai UR, et al. The alphaherpesvirus conserved pUS10 is important for natural infection and its expression is regulated by the conserved Herpesviridae protein kinase (CHPK). PLoS Pathog. 2023;19(2):e1010959. Epub 20230207. doi: 10.1371/journal.ppat.1010959.

58. Vega-Rodriguez W, Ponnuraj N, Jarosinski KW. Marek’s disease alphaherpesvirus (MDV) RLORF4 is not required for expression of glycoprotein C and interindividual spread. Virology. 2019;534:108–13. Epub 20190615. doi: 10.1016/j.virol.2019.06.008.

59. Ponnuraj N, Tien YT, Vega-Rodriguez W, Krieter A, Jarosinski KW. The Herpesviridae Conserved Multifunctional Infected-Cell Protein 27 (ICP27) Is Important but Not Required for Replication and Oncogenicity of Marek’s Disease Alphaherpesvirus. J Virol. 2019;93(4):e01903–18. Epub 20190205. doi: 10.1128/JVI.01903-18.

60. Krieter A, Ponnuraj N, Jarosinski KW. Expression of the Conserved Herpesvirus Protein Kinase (CHPK) of Marek’s Disease Alphaherpesvirus in the Skin Reveals a Mechanistic Importance for CHPK during Interindividual Spread in Chickens. J Virol. 2020;94(5):e01522–19. Epub 20200214. doi: 10.1128/JVI.01522-19.

61. Vega-Rodriguez W, Xu H, Ponnuraj N, Akbar H, Kim T, Jarosinski KW. The requirement of glycoprotein C (gC) for interindividual spread is a conserved function of gC for avian herpesviruses. Scientific reports. 2021;11(1):7753. Epub 20210408. doi: 10.1038/s41598-021-87400-x.

62. Thorvaldsdottir H, Robinson JT, Mesirov JP. Integrative Genomics Viewer (IGV): high-performance genomics data visualization and exploration. Brief Bioinform. 2013;14(2):178–92. Epub 20120419. doi: 10.1093/bib/bbs017.

63. Robinson JT, Thorvaldsdottir H, Turner D, Mesirov JP. igv.js: an embeddable JavaScript implementation of the Integrative Genomics Viewer (IGV). Bioinformatics. 2023;39(1). doi: 10.1093/bioinformatics/btac830.

64. Kim D, Paggi JM, Park C, Bennett C, Salzberg SL. Graph-based genome alignment and genotyping with HISAT2 and HISAT-genotype. Nat Biotechnol. 2019;37(8):907–15. Epub 20190802. doi: 10.1038/s41587-019-0201-4.

65. Wang L, Wang S, Li W. RSeQC: quality control of RNA-seq experiments. Bioinformatics. 2012;28(16):2184–5. Epub 20120627. doi: 10.1093/bioinformatics/bts356.

66. Li H, Handsaker B, Wysoker A, Fennell T, Ruan J, Homer N, et al. The Sequence Alignment/Map format and SAMtools. Bioinformatics. 2009;25(16):2078–9. Epub 20090608. doi: 10.1093/bioinformatics/btp352.

67. Cotto KC, Feng YY, Ramu A, Skidmore ZL, Kunisaki J, Richters M, et al. RegTools: Integrated analysis of genomic and transcriptomic data for the discovery of splicing variants in cancer. BioRxiv. doi: 10.1101/436634.

68. Quinlan AR, Hall IM. BEDTools: a flexible suite of utilities for comparing genomic features. Bioinformatics. 2010;26(6):841–2. Epub 20100128. doi: 10.1093/bioinformatics/btq033.

69. Benaglia T, Chauveau D, Hunter DR, Young DS. mixtools: An R Package for Analyzing Mixture Models. Journal of Statistical Software. 2009;32(6):1–29. doi: 10.18637/jss.v032.i06.

70. Kulak NA, Pichler G, Paron I, Nagaraj N, Mann M. Minimal, encapsulated proteomic-sample processing applied to copy-number estimation in eukaryotic cells. Nature methods. 2014;11(3):319–24. Epub 20140202. doi: 10.1038/nmeth.2834.

71. Rappsilber J, Mann M, Ishihama Y. Protocol for micro-purification, enrichment, pre-fractionation and storage of peptides for proteomics using StageTips. Nature protocols. 2007;2(8):1896–906. doi: 10.1038/nprot.2007.261.

72. Eng JK, Jahan TA, Hoopmann MR. Comet: an open-source MS/MS sequence database search tool. Proteomics. 2013;13(1):22–4. Epub 20121204. doi: 10.1002/pmic.201200439.

73. Kim S, Pevzner PA. MS-GF+ makes progress towards a universal database search tool for proteomics. Nat Commun. 2014;5:5277. Epub 20141031. doi: 10.1038/ncomms6277.

74. The M, MacCoss MJ, Noble WS, Kall L. Fast and Accurate Protein False Discovery Rates on Large-Scale Proteomics Data Sets with Percolator 3.0. Journal of the American Society for Mass Spectrometry. 2016;27(11):1719–27. Epub 20160829. doi: 10.1007/s13361-016-1460-7.

75. McIlwain S, Tamura K, Kertesz-Farkas A, Grant CE, Diament B, Frewen B, et al. Crux: rapid open source protein tandem mass spectrometry analysis. J Proteome Res. 2014;13(10):4488–91. Epub 20140909. doi: 10.1021/pr500741y.

76. Millikin RJ, Solntsev SK, Shortreed MR, Smith LM. Ultrafast Peptide Label-Free Quantification with FlashLFQ. J Proteome Res. 2018;17(1):386–91. Epub 20171108. doi: 10.1021/acs.jproteome.7b00608.

77. Team RC. R: A language and environment for statistical computing Vienna, Austria: R Foundation for Statistical Computing; 2022. Available from: https://www.r-project.org/.

78. Gatto L, Lilley KS. MSnbase-an R/Bioconductor package for isobaric tagged mass spectrometry data visualization, processing and quantitation. Bioinformatics. 2012;28(2):288–9. Epub 20111122. doi: 10.1093/bioinformatics/btr645.

79. Rost HL, Sachsenberg T, Aiche S, Bielow C, Weisser H, Aicheler F, et al. OpenMS: a flexible open-source software platform for mass spectrometry data analysis. Nature methods. 2016;13(9):741–8. doi: 10.1038/nmeth.3959.

80. Teufel F, Almagro Armenteros JJ, Johansen AR, Gislason MH, Pihl SI, Tsirigos KD, et al. SignalP 6.0 predicts all five types of signal peptides using protein language models. Nat Biotechnol. 2022. Epub 20220103. doi: 10.1038/s41587-021-01156-3.

81. Hallgren J, Tsirigos KD, Pedersen MD, Armenteros JJA, Marcatili P, Nielsen H, et al. DeepTMHMM predicts alpha and beta transmembrane proteins using deep neural networks. bioRxiv. 2022. doi: 10.1101/2022.04.08.487609.

82. Pagni M, Ioannidis V, Cerutti L, Zahn-Zabal M, Jongeneel CV, Hau J, et al. MyHits: improvements to an interactive resource for analyzing protein sequences. Nucleic Acids Res. 2007;35(Web Server issue):W433–7. Epub 20070601. doi: 10.1093/nar/gkm352.

83. Reese MG, Eeckman FH, Kulp D, Haussler D. Improved splice site detection in Genie. J Comput Biol. 1997;4(3):311–23. doi: 10.1089/cmb.1997.4.311.

